# Distinct Microbial Communities Within and On Seep Carbonates Support Long-term Anaerobic Oxidation of Methane and Novel pMMO Diversity

**DOI:** 10.1101/2025.02.04.636526

**Authors:** Magdalena J. Mayr, Sergio A. Parra, Stephanie A. Connon, Aditi K. Narayanan, Ranjani Murali, Antoine Crémière, Victoria J. Orphan

**Affiliations:** Division of Biology and Biological Engineering, California Institute of Technology, Pasadena, California, USA; Division of Geological and Planetary Sciences, California Institute of Technology, Pasadena, California, USA; School of Life Sciences, University of Nevada Las Vegas, Las Vegas, Nevada, USA; Geo-Ocean, UMR 6538 CNRS-Ifremer-UBO-UBS, Plouzané, France

**Keywords:** methane-derived authigenic carbonate (MDAC), endolithic, pmoCAB, autoendolith, chemoherm, resilience

## Abstract

At methane seeps worldwide, syntrophic anaerobic methane-oxidizing archaea and sulfate-reducing bacteria (ANME-SRB) promote carbonate precipitation and rock formation, acting as methane and carbon sink. While maintenance of active anaerobic oxidation of methane (AOM) within seep carbonates has been documented, the ANME-SRB reactivity to methane exposure remains uncertain. Surface-associated microbes may metabolize AOM-derived sulfide, maintain carbonate anoxia, and contribute to carbonate dissolution and higher trophic levels; however, these microbial communities are poorly described thus far. Here we provide new insights into microbial diversity, metabolic potential, activity, and resiliency within and on Southern Californian methane seep carbonates, by combining 16S rRNA and metagenomic sequencing, laboratory incubations, and BONCAT-FISH. *Ca*. Methanophaga (ANME-1) dominated the carbonate interiors across different seepage activities, based on sequencing, while the dominant SRB was *Ca*. Desulfaltia, potentially a new ANME partner. BONCAT-FISH revealed differences in ANME-1 cell activity, suggesting cell dormancy or DNA preservation at less active seep sites. Carbonate incubations from low activity seeps (≥24 months) showed an exponential AOM reactivation (44-day doubling time), suggesting seep carbonates remain potential methane sinks over dynamic seepage conditions. The surface-associated communities were distinct from the carbonate interior and other seep habitats, and highly heterogeneous. Surface ANME-SRB biofilms and sulfide-oxidizing bacterial mats were associated with high and intermediate AOM carbonates, potentially influencing carbonate precipitation/dissolution. Carbonate surfaces shared diverse aerobic methanotrophs with invertebrates, potentially serving as pool for animal epibionts. Besides particulate methane monooxygenases from aerobic methanotrophs, we found divergent forms including within a Methylophagaceae (GCA-002733105) MAG suggesting a new function within Methylophagaceae.

## Introduction

At methane seeps across the world’s oceans, anaerobic methanotrophic archaea (ANME) and sulfate-reducing bacteria (SRB) promote carbonate precipitation by producing alkalinity through bicarbonate and sulfide [1]. These carbonates form over hundreds to thousands of years [2–4] and persist even longer [5] storing carbon long-term [6] and serving as a rare deep-sea hardground surface for diverse animal communities [7, 8]. Methane seepage is dynamic over the long carbonate lifetime [9]. Sulfate-coupled anaerobic oxidation of methane (AOM) yields little energy and ANME-SRB doubling times are on the order of months [10, 11]. ANME-SRB entombed in carbonates and associated microbial signatures have been interpreted as fossilized remnants [12, 13]. Investigations have since demonstrated that carbonates host active ANME-SRB biomass, presumably in rock pores, capable of methane removal [14, 15]. However, cell activity levels in these carbonates have not been fully resolved and it is currently unknown if carbonates with low AOM activity can be reactivated upon reexposure to methane.

The carbonate-hosted microbial diversity and metabolic potential remains understudied, particularly compared to seep sediments. Previous studies mainly focused on the seep carbonate interior and commonly recovered ANME-1 (Methanospirareceae, synonym Methanophagaceae) [15–17]. The family ANME-1 comprises 14 recognized genera (gtdb R220), however, the lineages associated with seep carbonates require further investigation. Other taxa reported from seep carbonates include other ANME lineages (ANME-2ab, c), Atribacteria, Chloroflexi, Proteobacteria, and Desulfobacterota, including syntrophic Seep-SRB1 [14–16, 18], but the diversity of syntrophic Seep-SRBs present within seep carbonates remains to be assessed.

Surface-associated microbial communities on seafloor exposed carbonates reside at the water/carbonate interface, with potentially steep redox gradients, between oxygen in seawater, and sulfide generated from rock-hosted AOM. By consuming oxygen with sulfide or residual methane, these surface communities may impact local redox conditions and promote anoxic conditions within the rock interior [19]. Recently, experiments demonstrated that the acidity produced during aerobic methane and sulfide oxidation may dissolve seep carbonates [20, 21], highlighting the role of microorganisms in both their synthesis and dissolution. Additionally, seep carbonates host animal communities dependent on these rock-hosted chemosynthetic microorganisms [22], but focused studies on the diversity and metabolic potential of surface-associated microorganisms have been lacking.

Here we study microbes within and on deep-sea carbonates with varying AOM activity at two methane seep sites off of Southern California. Using a combination of 16S rRNA analysis, metagenomics, BONCAT-FISH, isotopic rate measurements, and long-term incubations our work highlights the diversity and ecophysiological breadth of methane-oxidizing and other chemosynthetic microorganisms living within and on the surfaces of seep carbonates. We further reveal resiliency of endolithic AOM communities to long-term fluctuations in methane supply, demonstrating the ability to reactivate sulfate-coupled methane oxidation and growth after months to years of minimal activity/dormancy.

## Methods

### Sampling

Carbonates were collected from two methane seep areas, Del Mar (1023 m depth) [23] and Santa Monica Mound (800 m depth) [24], off Southern California in May 2021 during the MBARI WF05-21 cruise on the R/V Western Flyer and ROV Doc Ricketts. Six samples, Rocks 1-4, 7 and 9, were collected from Del Mar. Further, two samples from Santa Monica Mound 800 (SMM800) with tube structure, Chimlet, and Protochimney, were included in this study. Where possible, sediment push cores and water samples adjacent to the carbonate samples were collected for 16S rRNA gene comparison (one water and sediment sample from Del Mar, three of each at SMM800). Carbonate samples were transferred into argon flushed mylar bags, submerged in nitrogen sparged 0.2 µm filtered Niskin bottom seawater from the site, and heat sealed for transport back to the laboratory for further analysis. The *in situ* orientation of these carbonates was reconstructed using video footage collected during the dive. In the laboratory we subsampled the rocks with a tile saw or drill press cleaned with ethanol and Nanopure water between samples. DNA samples were then stored at −80°C until further subsampling. Subsamples for cell counts were fixed 24 hours in 2% paraformaldehyde at 4°C, washed twice with 3xPBS (phosphate-buffered saline), and stored in 70% ethanol 30% 1xPBS at −20°C. Sample coordinates and a detailed processing description are provided in the supplementary methods.

### AOM rate measurements

We measured AOM rates using mono-deuterated methane (CH_3_D) or ^13^CH_4_ as substrates according to [25, 26], and quantified δD-H_2_O or ^13^C-CO_2_ production, respectively, at five timepoints. At the last timepoint (t4) after 48-143 days (further details supplementary table 1), we measured sulfide. The incubations with carbonate subsamples were performed in serum vials with site-specific seawater at 4°C, close to *in situ* temperatures, under anoxic conditions. We added 100% or 50% labeled mono-deuterated methane (CH_3_D), and 100% or 10% labeled ^13^C-CH_4_ at 2 bars partial pressure of methane to the respective incubations. For subsequent BONCAT-FISH we further added 200 µM HPG (L-Homopropargylglycine). We performed unlabeled and killed controls in parallel incubations. The incubations with ^13^CH_4_ did not yield a quantitative rate measurement for most samples. While we show the results in supplementary table 1, we do not discuss them further.

We derived the anaerobic methane activation rates from the slope of a linear regression of δD-H_2_O production over time, measured with a Picarro isotopic water analyzer (2140-i). Using CH_3_D may underestimate rates if not all four hydrogen atoms form water during methane oxidation, thus representing a conservative estimate. Still, CH_3_D is potentially more sensitive than ^13^CH_4_, because full oxidation to CO_2_ is not necessary, such that methane activation is enough for a signal. We therefore report the rates as anaerobic methane activation rates in nmol Deuterium per cm^−3^ d^−1^. The δD-H_2_O of the killed controls did not increase over time. For a more detailed discussion of CH_3_D rate measurements, see [25]. Incubations without measurable enrichment of δD-H_2_O over time are reported as rates below detection. Further details on the rate measurements are provided in the supplementary methods.

### Long-term reactivation incubations

For long-term reactivation incubations, the remaining rocks of R1-R4 (approx. 50 to 300 g) were incubated in Duran bottles with sterile, anoxic artificial seawater (approx. 100 – 400 mL, recipe see supplementary table 2) and a 2.4 bar unlabeled methane headspace. Sulfide was monitored over time and measured photometrically [30] on a plate reader (TECAN Sunrise). Further details are provided in the supplementary methods.

### DNA extraction

For DNA analysis the frozen rocks were further subsampled into 4-7 horizons in 1-7 mm steps from surface to interior. The rock surface was scraped off with an ethanol flamed spatula and the rock horizons were cut using a rotary tool (Dremel) with a diamond wheel and then ground to powder. Rocks and sediment were extracted with the DNeasy PowerSoil Pro Kit (Qiagen) with modifications for rocks similar to [27]. Water samples filtered onto Sterivex filters were extracted with a phenol-chloroform method. Further details are provided in the supplementary methods.

### 16S rRNA gene sequencing and analysis

The 16S rRNA gene (V4-V5) was amplified using 515F and 926R [28] archaeal/bacterial primers with Illumina adapters in duplicate PCR reactions with 33 cycles and 54 °C annealing temperature (Q5 Hot Start High-Fidelity 2x Master Mix, New England Biolabs, USA). Duplicates were pooled and barcoded with Illumina Nextera XT index 2 primers. Barcoded PCR products were combined equimolarly, purified, and sequenced on Illumina’s MiSeq platform with 15-20% PhiX by Laragen (Culver City, CA). Further details provided in the supplementary methods.

After sequencing we removed adapters with cutadapt (v. 3.4) and inferred amplicon sequence variants (ASVs) using DADA2 (v. 1.20.0) in R (v. 4.2.2). ASVs were annotated with IDTAXA of DECIPHER (v. 2.20.0) and the SILVA_SSU_r138 database amended with inhouse sequences. Decontam (v. 1.18.0) in R was used to remove contaminants. For a refined ANME-1 classification we extracted available 16S rRNA genes from gtdb representative genomes with barrnap (v0.9) [29]. Together with sequences from this study we constructed a phylogenetic tree using IQtree (v2.1.2) [30] (Fig. S4). The SILVA database currently only distinguishes ANME-1a and ANME-1b, thereby underestimating the ANME-1 taxonomic diversity of 14 genera present in gtdb. Further, the 16S rRNA gene trees of SRB (Fig. S5) and aerobic methanotrophs (Fig. S9) were constructed with IQtree. Sequences were aligned with muscle (v. 3.8.1551).

### Metagenomics sequencing and analysis

We selected three rocks, R1, R9 and Chimlet, for metagenomic sequencing of the interior and surface. We prepared the library with the Illumina DNA Prep kit and 10-12 amplification cycles depending on input. The Keck Genomics Platform of the University of California quantity- and quality-checked the libraries with a TapeStation 4200 (Agilent) and sequenced them on a NovaSeq 6000, S1 flowcell, for 150 bp paired-end reads (Illumina Inc.).

Using bbduk [31] we trimmed primers and adapters. We assembled the reads five times using a) megahit (v1.2.9) [32] coassembly b) metaspades (v3.15.2) [33] individual assemblies c) megahit individual assemblies d) megahit coassemblies normalized with bbnorm (target=33) [31], and e) megahit bbnorm individual assemblies. Further details provided in supplementary methods. We binned and refined mags using metawrap (v1.3.2, concoct, maxbin2, metabat2) [34] from each assembly with reads from all samples for differential coverage. We then dereplicated the bins at 95% ANI with dRep (v2.6.2) [35]. Selected bins were manually inspected with anvio 7.1 [36], and obvious contamination was removed based on evenness of coverage. Mdmcleaner [37] was used to taxonomically classify contigs and was used in addition to anvio visualization to guide manual refining in anvio 7.1 if necessary. We classified the mags with gtdbtk (v2.3.2, database version r220) [38] and determined MAG coverage with coverM (v0.6.1) [39]. The mags were annotated with metabolic (v4.0) [40]. Based on checkM and checkM2 [41, 42] MAGs with >50% completeness and <10% redundancy were kept. MAGs that dropped below 50% completeness after refinement were kept, prioritizing lower contamination over completeness.

Phylogenetic trees of ANME and SRB MAGs were done with anvio 7.1 using the Archaea_76 and Bacteria_71 marker genes set, respectively. Reference genomes were retrieved from gtdb r214 and amended with ANME-1c genomes [43]. We updated taxonomic names to gtdb r220 where necessary.

### *pmoC* analysis

The first set of divergent *pmoC* sequences was retrieved from annotating MAGs with metabolic. Only the bin_133 (Methylophagaceae) unambiguously contained the divergent *pmoC* based on anvio inspection. Further *pmoC* sequences were searched and extracted from translated predicted genes (prodigal v2.6.3) [44] from the metagenomic assemblies using diamond blastp (v2.0.6.) [45]. A *pmoC* database was constructed for this purpose. Reference sequences of cultivated and uncultivated microorganisms were either retrieved using NCBI blastp [46] or were extracted from MAGs from gtdb. The *pmoC* gene tree was constructed with IQtree. All trees were visualized with iTol (v.6) [47]. The *pmoC* sequences from this study are provided as a supplementary fasta file.

### Cell extraction

To extract cells from the carbonate rock matrix for microscopy, we developed an extraction protocol combining a cell extraction buffer [48] with a percoll density centrifugation [49]. A detailed protocol is available on protocols.io [50].

### Cell counts

Extracted cells were filtered onto black, 0.2 µm pore size polycarbonate membranes (GTBP02500, Isopore, Millipore Sigma), and mounted with Citifluor containing 4′,6-diamidino-2-phenylindole (DAPI, 4.5ng/µL). We analyzed between 13-30 field of views and 2320 - 11502 cells per sample on an Elyra PS.1 SIM microscope (Zeiss, Germany) with alpha Plan-APOCHROMAT 100X/1.46 oil objective. Images with lower quality were removed prior to automatic counting with Fiji-ImageJ (2.3.0, v. 1.53q, Java 1.8.0_172). Automatic counting included Gaussian blur 1 for noise reduction, background subtraction (rolling ball, 50 pixels) and thresholding to create masks (Otsu). Particles with a minimum size of 0.04 µm^2^ were counted after watershed transformation.

### Translational activity of ANME with BONCAT-FISH

Extracted cells from selected HPG-incubated rocks (R1, R3, R9, Chimlet, Protochimney) were mounted onto slides and HPG incorporation was visualized using a click-reaction with AF647 picoyl azide [51]. Fluorescence *in situ* hybridization (FISH) was done according to standard protocols [52], using an archaeal (ARCH915 [53], dualAlexa546) and a bacterial probe mix (EUB, EUBII and EUBIII [52], dualAlexa488). We interpret the archaeal cells in R9, Chimlet and Protochimney as coming almost exclusively from ANME, by far the most abundant archaea based on 16S rRNA gene amplicon sequencing and metagenomics. The Del Mar outcrop showed some other archaeal community members, but we did not find any archaeal cells (see results). We confirmed ANME-1 presence in R9. To do so we mixed ANME1-350 [54] (cy3, 5’-AGTTTTCGCGCCTGATGC-3’) and a new probe ANME1-728 (cy3, 5’-GGTCTGGTCAGACGCCTT-3’) designed in ARB using SILVA 138 database, both highly specific. Two probes targeting different regions on the 16S rRNA were combined to obtain a stronger signal. For data analysis we identified archaeal cells automatically and inspected the archaeal cells manually for a positive BONCAT signal, which was counted when showing a cell-shape in the cy5 (BONCAT) channel. Further details on BONCAT-FISH and analysis are provided in the supplementary methods.

## Results

### Seep carbonates with low to high AOM activity

Carbonates from the Del Mar and Santa Monica Mound 800 (SMM800) methane seeps off Southern California (Fig. 1a,b [55]), were characterized with *in situ* observations. The Del Mar outcrop (Rocks 1-4) extended into the water column (near lower edge of OMZ, 22 µM oxygen) and lacked active methane seepage and AOM indicators such as sulfide-oxidizing microbial mats and chemosynthetic animals (Fig. 1c), suggesting low AOM activity. Del Mar Rock 9 was covered in white sulfide-oxidizing bacteria (Fig. 1d), suggesting active AOM and sulfide production. Chimlet and Protochimney from SMM800 are hollow carbonate tubes sealed at the top and extended into an OMZ (8 µM oxygen). White and black microbial mats, and animals covered these rocks suggesting high AOM activity (Fig. 1e,f). Relative to Chimlet and Protochimney, we refer to Rock 9 as having intermediate AOM activity. Depleted δ^13^C_carbonate_ values (−46.2 to −54.0‰) confirm the methane-derived origin [56] of the studied carbonates (Fig. S1).

**Figure 1.**
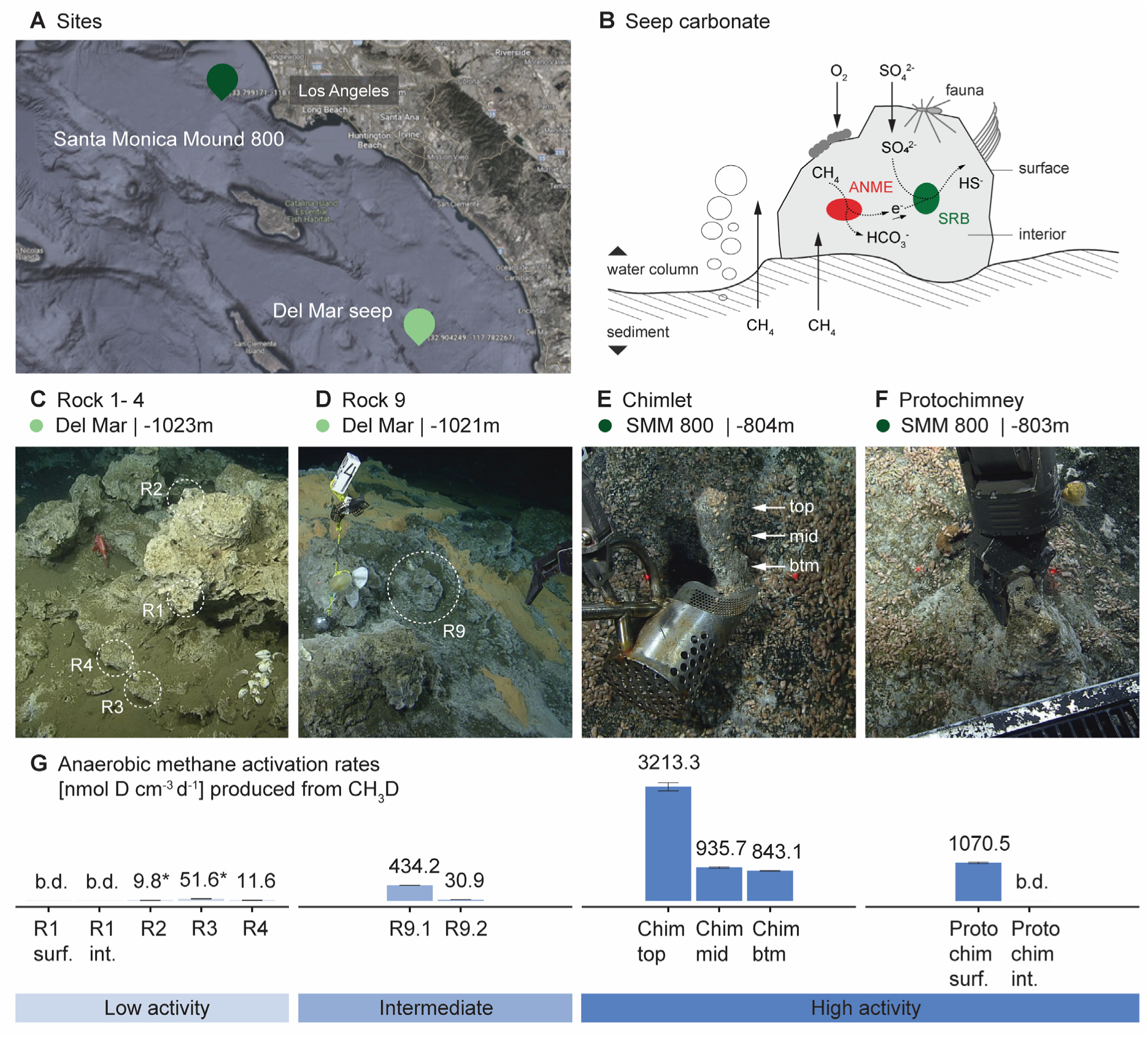
Anaerobic methane oxidation activity of methane seep carbonates from Del Mar and SMM800. *In situ* AOM indicators and CH_3_D rate measurements characterize low to high AOM activity carbonates. **A**) Seep carbonate collection sites Del Mar (light green marker) and Santa Monica Mound 800 (SMM800, dark green marker) are located 129 km apart. Map obtained from Google Maps. **B**) Biogeochemical seep carbonate setting. *In situ* images of **C**) the Del Mar outcrop, R1 and R2 originated from the top, R3 and R4 from closer to the sediment. **D**) R9, from a nearby Del Mar area with sulfide oxidizing mats. **E**) Chimlet and **F**) Protochimney are two chemoherm-like structures and were collected from different sides of the Santa Monica Mound 800. Chimlet actively bubbled with methane upon recovery. For scale, the red laser points in the images are 29 cm apart. **G**) Anaerobic methane activation rates (nmol D cm^−3^ d^−1^) measured in anoxic incubations with monodeuterated methane based on: CH_3_D + SO_4_^2-^ ◊ HCO_3_^−^ + HS^−^ + HDO. We measured δD of water over five timepoints and calculated the rate from a linear increase unless stated otherwise. Error bars show the standard error of k calculated from the linear regression. Two subsamples of R9, R9.1 and R9.2 with different color, light grey and dark grey, respectively, were incubated for AOM rates. The orientation of the R9 piece dedicated for rates could not be reconstructed. At the last time point (t4) sulfide was measured and was detectable in R9.1, Chimlet top, middle, bottom, and Protochimney surface. *Deuterium above background was only detected at t4 indicating a nonlinear increase in R2 and R3., b.d. below detection, surf. surface, int. interior, btm. bottom

We tested the AOM potential using anoxic incubations amended with monodeuterated methane (CH_3_D), and measured methane activation via HDO production [25] over 1.6 to 4.8 months. The Del Mar outcrop carbonates, Rock 1-4, had low anaerobic methane activation rates, from non-detectable to 11.6 nmol D cm^−3^ d^−1^, without measurable sulfide production within 4+ months. Of these, R2 and R3 showed an HDO increase at the last time point (4+ months), potentially indicating stimulation of ANME-SRB growth (Fig. 1g). Based on observed color differences two Rock 9 subsamples, R9.1 (light grey) and R9.2 (dark grey) were incubated. R9.1 showed anaerobic methane activation and sulfide production, confirming *in situ* indicators of active sulfate-coupled AOM. R9.2 had a lower anaerobic methane activation potential. Differences between subsamples are expected, as carbonate rocks are naturally heterogeneous. The tubular Chimlet sample had high anaerobic methane activation rates, with the greatest rates associated with the cap (Fig. 1e,g). The outer surface sample of Protochimney also showed high anaerobic methane activation rates, while the corresponding interior piece displayed no measurable activity. This likely does not reflect total inactivity in the interior, as a second Protochimney interior piece showed sulfide production (data not shown). Combined with *in situ* observations and the high AOM potential, we refer to Chimlet and Protochimney, both from SMM800, as high AOM activity carbonates.

### Distinct seep microbial communities – heterogeneous carbonate surface and homogeneous interior

Rock subsectioning and 16S rRNA gene sequencing revealed a thin, heterogeneous surface microbial community and a more homogeneous interior community beneath it, both distinct from other seep habitats (Fig. 2a,b). The surface veneer community (0-1mm) was different from the interior and comparatively heterogenous (Fig. 2a,b), often showing high relative abundances of Proteobacteria (e.g. unclassified Gammaproteobacteria, Methylococcales, Chromatiales, Thiotrichales, Beggiatoales) in contrast to the interior (Fig. 2c). The 1-2mm section included the previously scraped surface, sometimes with surface community remnants (Fig. 2c, e.g. R1). The Chimlet and Protochimney surfaces were dominated by ANME-SRB (Halobacteriota, Desulfobacterota), like the interior at phylum level. Methanocomedenaceae and Seep-SRB1a dominated these biofilms, and on Protochimney additionally *Ca*. Methanogaster and Seep-SRB2 (Fig. 2d,e). White/orange surface regions, again, showed more Proteobacteria (Fig. 2c). The interior hosted high relative abundances of Halobacteriota (primarily ANME), and variable abundances of Caldatribacteriota, Desulfobacterota, Planctomycetota and Asgardarchaeota (Fig. 2c, additional profiles Fig. S2, median abundance ranks Fig. S3). The surface microbial community was distinct from the water column (Fig. 2b), even though they are in contact. Both rock and sediment surfaces are redox and phase transition zones, however, their communities were clearly distinct (Fig. 2b).

**Figure 2.**
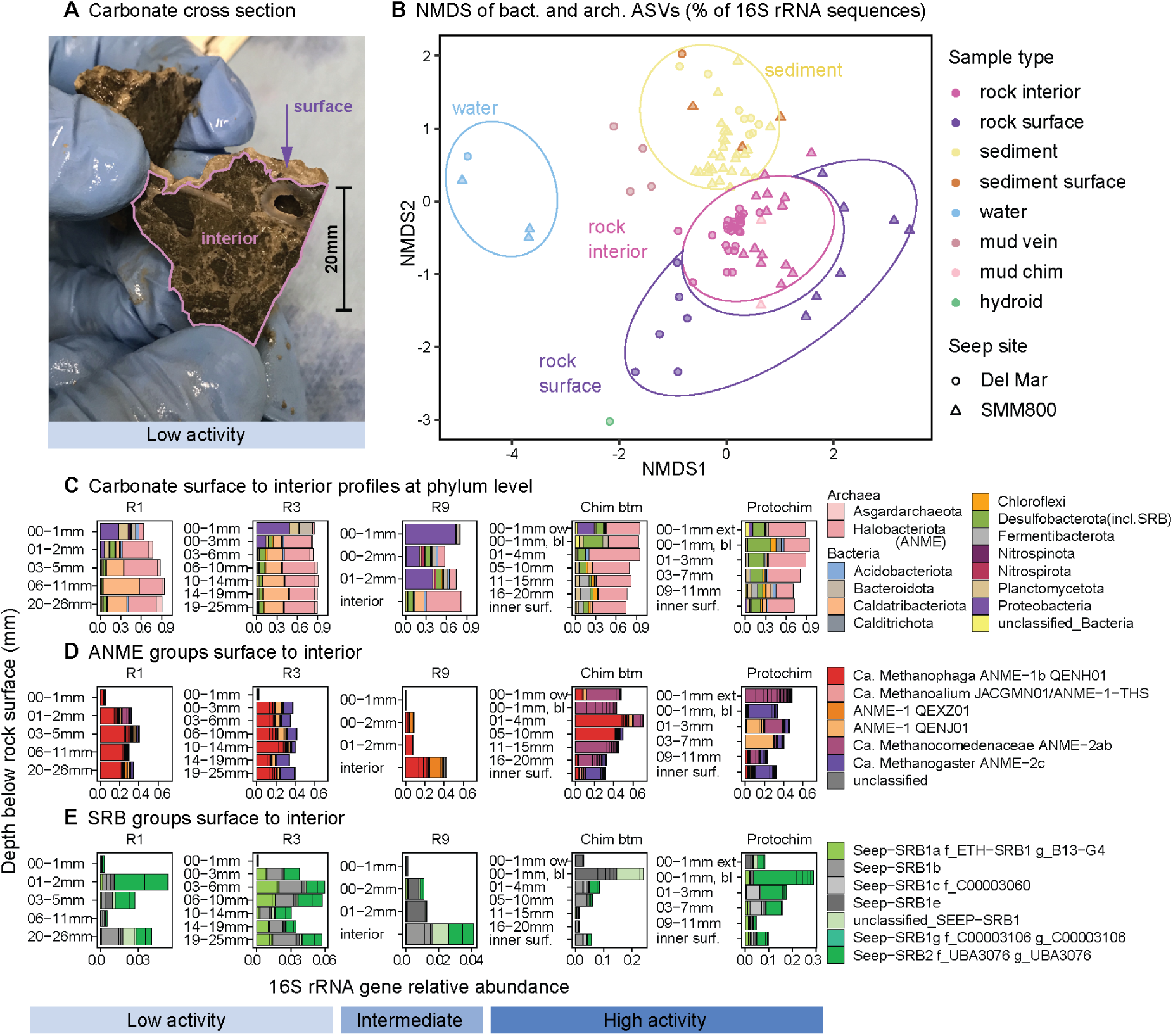
Microbial communities from the carbonate surface to interior in context of other the seep communities. 16S rRNA gene sequencing of carbonate surface scrapes (0-1mm) and mm-scale sections. **A**) Cross section of Del Mar outcrop Rock 4, with a white partially scrapeable surface layer (0-1mm). Chimlet and Protochimney had a black biofilm instead with white and orange patches. **B**) NMDS of bacterial and archaeal ASVs showing dissimilarities between rock surface and interior in context of sediment, water and other seep communities. Each point represents one sample. (n=103, stress=0.18). Shapes were hand drawn for visualization purposes. **C**) Archaeal and bacterial phyla in selected sectioned carbonates from surface to interior based on 16S rRNA gene sequencing. Phyla reaching ≥5% at least once are shown. Carbonates naturally varied in size and shape which led to different section sizes, even though we tried to cut them as similarly as possible. We sectioned three parts of Chimlet: top (cap), middle, bottom (btm). **D**) ANME phylogenetic groups based on gtdb taxonomy identified with a 16S rRNA phylogenetic tree (Fig. S4) to genus level where possible. **E**) SRB phylogenetic groups based a 16S rRNA phylogenetic tree (Fig. S5). Note that SRB x-axes are scaled to the maximum abundance of the respective profile for better visibility, because SRB abundances varied substantially. ANME and SRB ASVs reaching ≥1% at least once were included. See full set of sectioned rock microbial communities in supplementary Fig. S2 (including R2, R4, R7, Chimlet top, Chimlet mid). Abbreviations: Inner surf., inner surface of Chimlet and Protochimney cavity; bl, black; ow, orange white

We found mm-scale open veins with sediment-like material in the Del Mar outcrop, potentially inhabited by animals, with microbial communities between that of water column, sediment, and rock (“mud vein”, Fig. 2b). Mud recovered from the central cavity of Chimlet harbored a microbial community that was similar to the lithified rock (“mud chim”, Fig. 2b). Del Mar rock R7, had an attached hydroid whose microbiome was most similar to, but distinct from the Del Mar carbonate surface communities.

### Dominant endolithic ANME-1 genera and a potential new SRB partner

Based on 16S rRNA gene sequencing and metagenomics, the ANME-1 genus *Ca*. Methanophaga (QENH01, ANME-1b) dominated the rock interior, and the dominant SRB MAG was *Ca*. Desulfaltia. We recovered further ANME-1 MAGs affiliated with the genera QEXZ01 and QENJ01, that were also present in the rock interiors (Fig. 3). Within Chimlet and Protochimney, we identified JACGMN01 (*Ca*. Methanoalium-2, ASV_537, tree Fig. S4) based on 16S rRNA amplicons at low relative abundance, totaling to 3-4 ANME-1 genera per rock interior. Notably, high ANME-1 relative abundances were observed independent of the AOM activity measured (Fig. 2d, 3).

**Figure 3.**
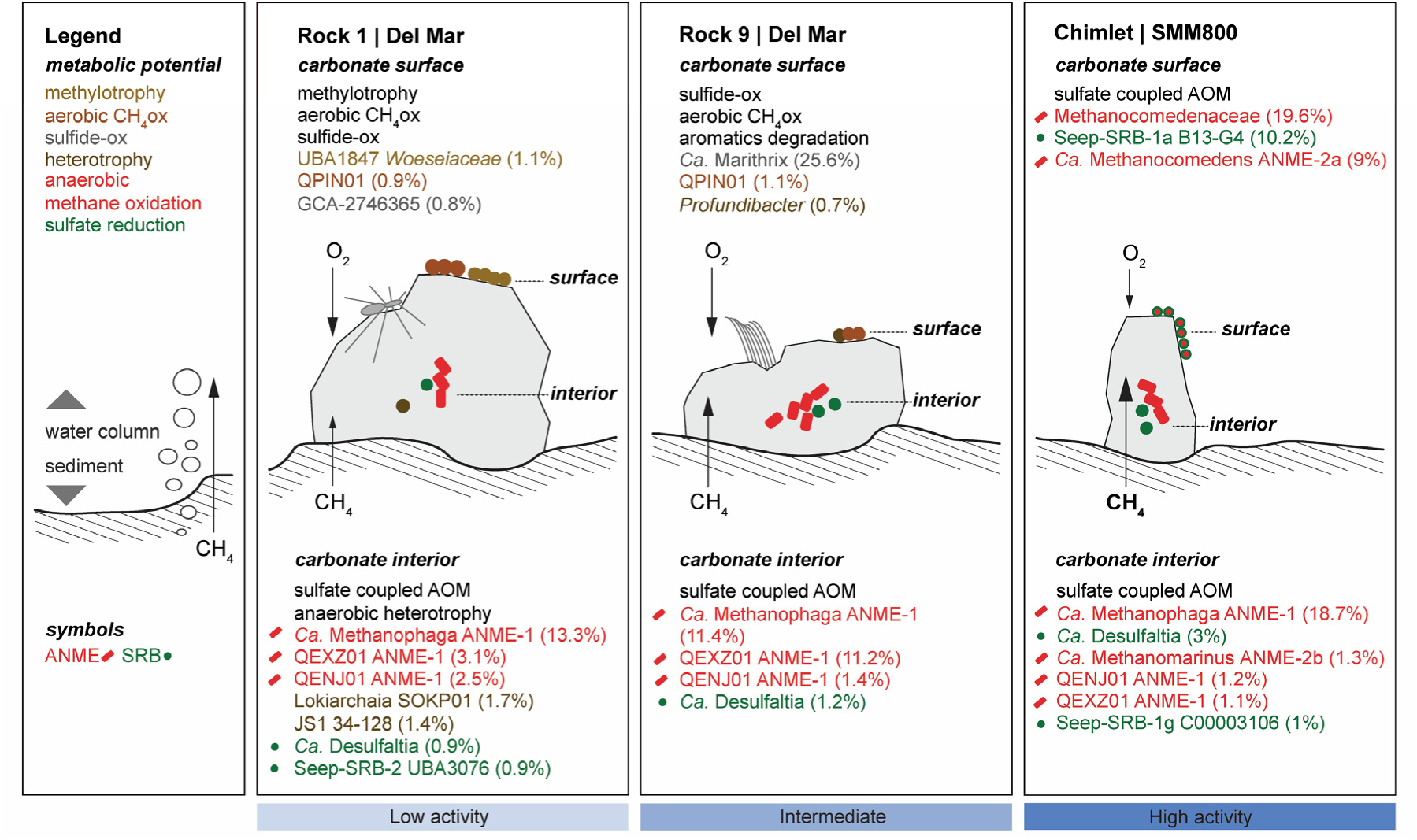
MAGs and metabolic potential in and on carbonates with low, intermediate and high AOM activity. We sequenced the surface and the interior of R1, R9 and Chimlet with low, intermediate and high AOM activity, respectively, and binned MAGs from these six metagenomes to investigate taxonomy and metabolic potential. MAGs shown were selected based on relative abundance, shown in parenthesis, with a focus on *Proteobacteria* on the rock surface because of their higher abundance compared to the interior (Fig. 2c). Find a full list of MAGs in supplementary table 3. Anaerobic methane oxidizing archaea dominated the rock interiors, *Ca*. Methanophaga, in particular, and *Ca*. Desulfaltia was the most abundant interior SRB suggesting it may be an ANME partner bacterium. In contrast, the surface communities and their metabolic potentials differed between rocks along with the AOM activity of the carbonate, methane supply strength, and oxygen concentrations. We indicated the difference in methane supply to the rocks estimated based on the *in situ* observations with arrow size. Further, oxygen concentrations were higher at the Del Mar (22 µM) vs. SMM800 (8 µM) seep.

In contrast, Chimlet’s surface mat was dominated by Methanocomedenaceae (ANME-2ab). Additionally, a few rock sections adjacent to the inner cavity walls of Chimlet and Protochimney showed the presence of Methanocomedenaceae (Fig. 2d). Members of *Ca*. Methanogaster (ANME-2c) were additionally detected in the Del Mar outcrop carbonate interiors and dominant in the inner cavity walls of Chimlet and Protochimney and the Protochimney biofilm (Fig. 2d).

We recovered MAGs of known ANME partner bacteria including Seep-SRB-1a (B13-G4), Seep-SRB-1g (C00003106), and Seep-SRB-2 (UBA3076). However, *Ca*. Desulfaltia (bin_051, 65% completeness) was the dominant interior SRB MAG (Fig. 3). *Ca*. Desulfaltia belongs to the family ETH-SRB1, which contains Seep-SRB-1a, and the *Ca*. Ethanoperedens-partner genus ETH-SRB1 (Fig. 4b). Bin_051 lacked a 16S rRNA gene, but based on gtdb, *Ca*. Desulfaltia may belong to Seep-SRB-1b [57]. Large multiheme cytochromes (*oetAB*) predicted to be involved in the electron transfer from ANME to SRB [58] were missing from bin_051, but were detected in two *Ca*. Desulfaltia gtdb MAGs (Fig. S6). Based on these results, *Ca*. Desulfaltia is a promising candidate for a syntrophic ANME partner bacterium, but additional evidence is required.

**Figure 4.**
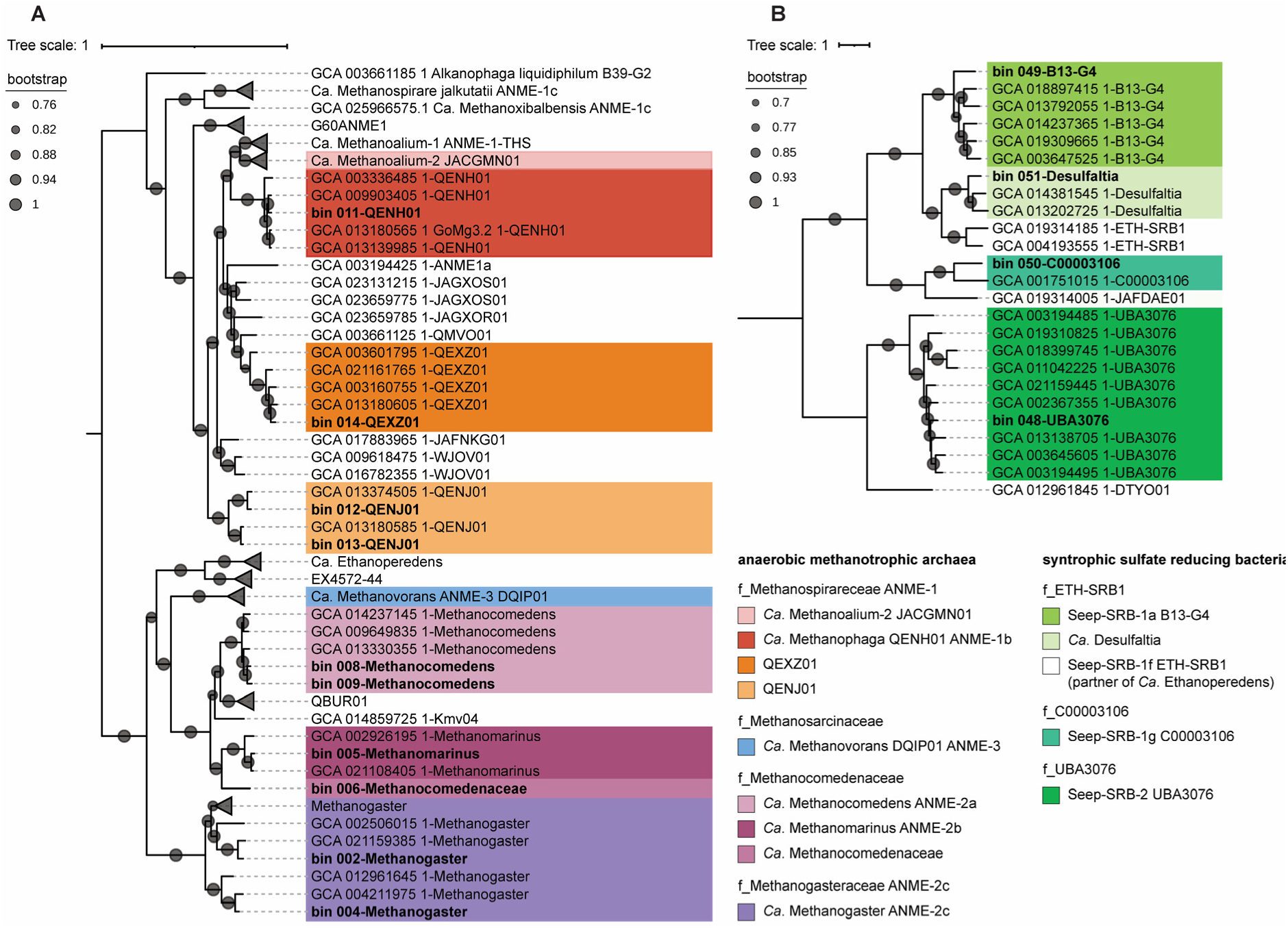
Phylogenomic tree of ANME and Seep-SRB metagenome assembled genomes (MAGs) and reference MAGs. MAGs from this study are in bold. We give the current gtdb taxonomy, previously proposed names, and historic names for reference. **A**) We find four ANME-1 MAGs within three genera with highest relative abundance in the rock interior. *Ca*. Methanocomedens and bin_006 Methanocomedenaceae, which may be part of a new genus had highest abundance in Chimlet’s black biofilm. *Ca*. Methanomarinus was present at lower relative abundance in Chimlet’s interior. We found two *Ca*. Methanogaster MAGs, reaching a max. of 0.6% rel. ab. in Chimlet’s interior. **B**) We found MAGs of the known ANME-partner genera Seep-SRB-1a, Seep-SRB-1g and Seep-SRB-2, with highest rel. ab. in Chimlet’s surface, Chimlet’s interior, R1’s interior, respectively (Fig. 3). The most abundant SRB MAG in the rock interior was *Ca*. Desulfaltia, an uncultivated genus part of family ETH-SRB1. ETH-SRB1further contains Seep-SRB-1a, a known ANME partner, and genus ETH-SRB1, a known sulfate reducing partner bacterium of *Ca*. Ethanoperedens.

### Investigating the mismatch between ANME relative abundance and AOM activity with BONCAT-FISH

Strikingly, carbonates R1-R4 with low AOM activity, showed high DNA-based ANME relative abundances (Fig. 1, 2d, 3). Further, both low and high AOM activity rocks had similar cell counts (8×10^7^-7×10^8^ g^−1^, Fig. 5a), suggesting that the active cell proportions varied. This scenario was tested with BONCAT-FISH after anoxic incubation with methane for approx. 50-140 days. We used an archaeal FISH probe to stain ANME, because most archaeal sequences were affiliated with ANME in the tested rocks (Fig. 2c). The higher AOM activity rocks R9, Protochimney and Chimlet had a large fraction of BONCAT-positive ANME cells representing 41%, 68% and 83%, respectively. Many of the cells displayed the commonly described ANME-1 rectangular rod shape [49] (Fig. 5b,c). Using R9, we confirmed abundance of ANME-1 using group-specific FISH probes (representative image, Fig. S7). This contrasted with the low AOM activity rocks R1 and R3, where we could not identify archaeal cells by FISH, and, therefore, no BONCAT-active archaea either, which may indicate ANME cells were dormant or low activity, or active cells were too rare to be detected.

**Figure 5.**
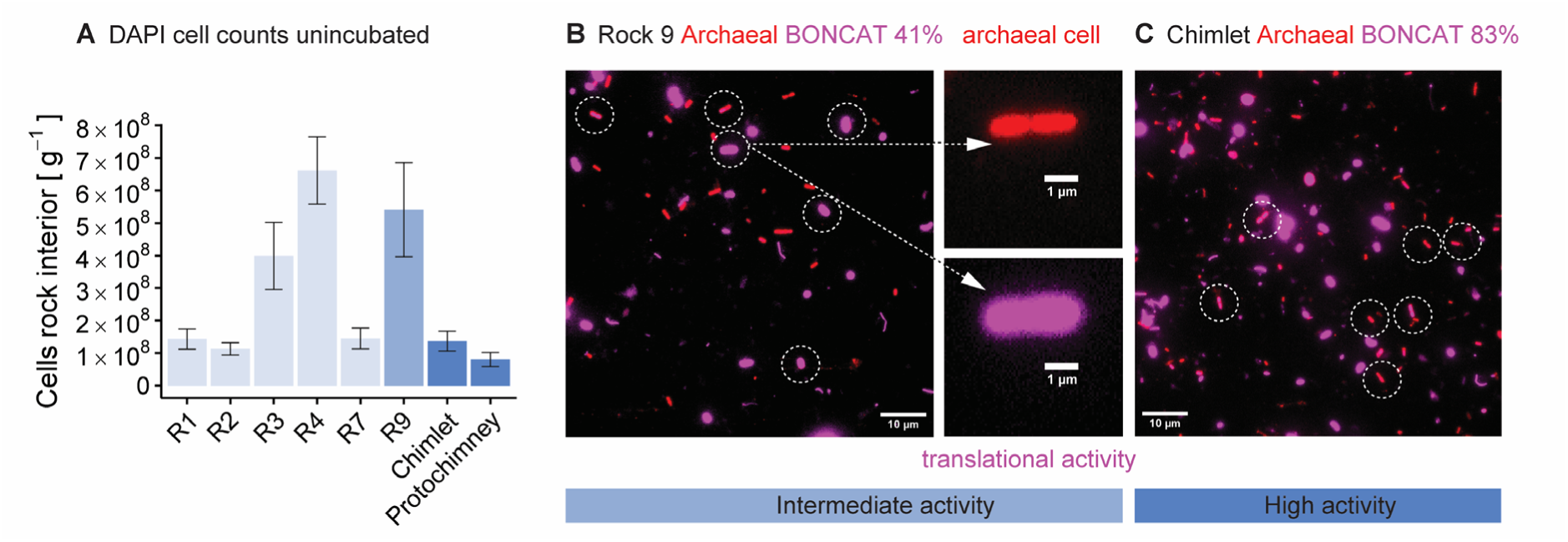
Cell counts of carbonate-hosted microorganisms and BONCAT-FISH of carbonate-hosted ANME. **A**) DAPI cell counts of extracted cells from the carbonate interior pre-incubation. Error bars represent the standard deviation between different fields of view. Representative BONCAT-FISH images of **B**) Rock 9 and **C**) Chimlet. BONCAT-FISH was used to examine the anabolic activity of endolithic ANME cells after anoxic incubation with methane and HPG for approx. 50-140 days. Bioorthogonal noncanonical amino acid tagging (BONCAT) measures the translational activity at the single cell level via incorporation of HPG, a methionine analog. Using fluorescence *in situ* hybridization (FISH) we identified archaeal cells (red, ARCH915). White circles highlight examples of archaeal cells (red) with corresponding BONCAT signal (magenta). BONCAT intensities varied substantially between cells. Arrows in **B**) point to magnified representative cells for which archaeal FISH signal (red, upper image) with typical ANME-1 shape and BONCAT signal (magenta, lower image) are shown separately. R9, Protochimney, and Chimlet had 41%, 68%, and 83% BONCAT positive archaeal cells, respectively.

### Reactivation of carbonates with low AOM in long-term incubations

To test if low AOM activity carbonates have the capacity to regain AOM activity, we conducted long-term incubations (700-1000 days) under conditions mimicking methane reactivation. After a lag time of about 300 days, the sulfide concentration of R4 increased exponentially, suggestive of ANME-SRB growth [59, 60] with an estimated rate (r) of 0.0157 d^−1^ and apparent doubling time of 44 days (Fig. 6a). No electron donors besides methane were added. Similarly, adjacent R3 and R1 carbonates showed exponential sulfide production after approx. 600 and 700 days, respectively (Fig. 6b,c). The growth parameters of R2 and R3 should be treated with caution because of lower R^2^ and sampling frequency. Our results suggest AOM can be restored over months to years once methane becomes available.

**Figure 6.**
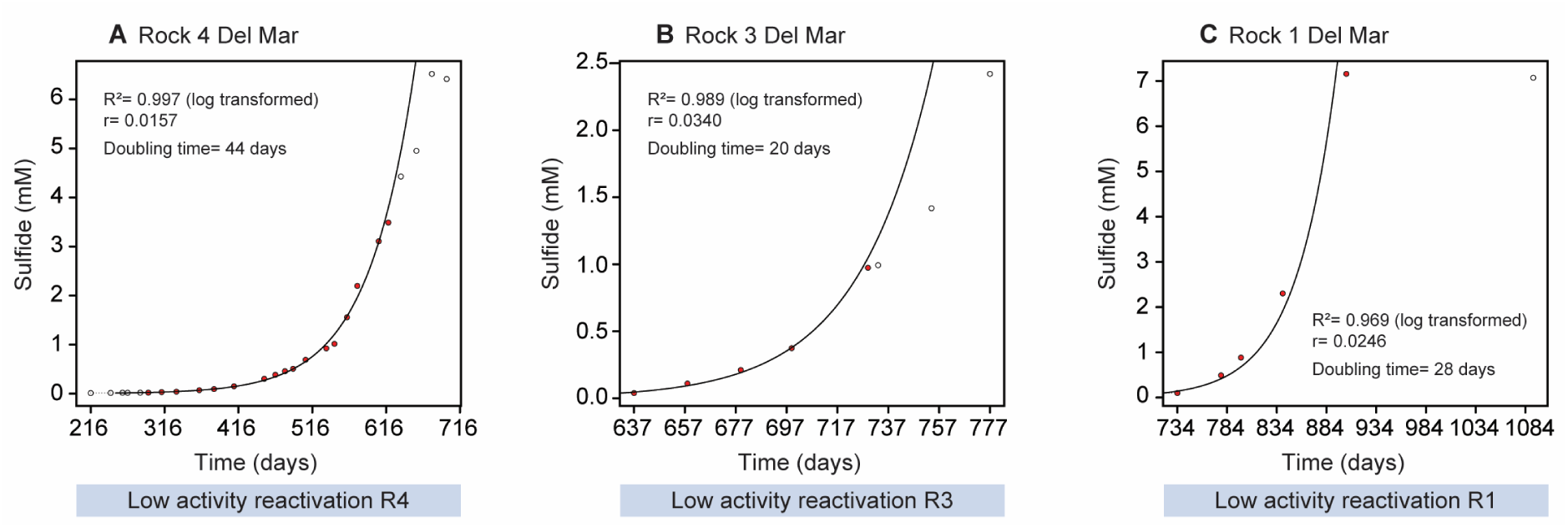
Exponential sulfide increase in long-term incubations of low AOM carbonates with sulfate and methane. We reactivated the initially low AOM activity carbonates **A**) R4, **B**) R3 and **C**) R1, in order of reactivation, in long-term incubations mimicking methane resurge with artificial seawater (10 mM sulfate) and a methane headspace under anoxic conditions. The incubation of carbonate R2 was accidentally lost. The red dots (n=17, n=5, n=5, respectively) were included in the growth rate calculation (for details see methods). Note that only the phase with increasing sulfide concentrations is shown. The total monitoring time was 698, 777, and 1092 days, respectively. R^2^ and growth rate r were derived from a linear regression of log transformed concentrations (Fig. S8). The observed lag time of months to years might be explained by few initial viable ANME-SRB or a long growth preparation time. The slowing of the exponential increase at the last timepoints might point to a resource limitation.

### Diversity and metabolic potential of the surface community reflects carbonate-associated AOM activity

Proteobacteria dominated the surface of the low AOM carbonate R1. No single MAG dominated, and gammaproteobacterial bin_147 (UBA1847, Woeseiaceae) had the highest relative abundance with 1.1% (Fig. 3). This MAG contained genes for sulfur oxidation, methanol and methanethiol utilization, nitrite reduction and autotrophy (form II Rubisco). The most abundant aerobic methanotroph, bin_129 (QPIN01, Methylomonadaceae, 0.9% rel. abundance), encoded nitrite reduction and sulfide oxidation genes. Gammaproteobacterial bin_139 (GCA-2746365, SZUA-229, 0.8% rel. abundance) contained genes for sulfide oxidation, carbon fixation (form I Rubisco) and nitrite reduction. Gammaproteobacterial bin_124 (GCA-001735895, 0.6% rel. abundance) encoded genes for methanol and methanethiol utilization, carbon fixation (form I Rubisco), nitrate and nitrite reduction, and sulfide oxidation. Supplementary table 3 contains a complete list of MAGs.

The surface of carbonate R9 with higher AOM activity rates was dominated by large sulfur-oxidizing *Ca*. Marithrix (bin_119, Beggiatoaceae, 25.6% rel. abundance, Fig. 3), consistent with the white *in situ* mat (Fig. 1). The MAG encoded nitrate, nitrite, and nitric oxide reduction genes as previously described [61], and likely utilizes sulfide generated by endolithic ANME-SRB. Like R1, R9 again hosted the aerobic methanotroph genus QPIN01 (bin_130, 1.1% rel. abundance). Further, Alphaproteobacteria bin_112 (*Profundibacter*, Rhodobacteraceae, 0.7% rel. abundance) encoded genes for benzoyl-coA reduction, formaldehyde utilization, nitrate reduction to N_2_, sulfide oxidation and carbon fixation (form II Rubisco).

Chimlet from SMM800 had a distinct surface community from the Del Mar surface communities, here dominated by an ANME-SRB biofilm. Chimlet’s biofilm was dominated by ANME-2 bin_006 (Methanocomedenaceae, 19.6% rel. abundance, Fig. 3), a potentially new Methanocomedenaceae genus (Fig. 4a), followed by the ANME partner bacterium Seep-SRB-1a (bin_049, 10.1% rel. abundance), and bin_009 belonging to *Ca*. Methanocomedens (9% rel. abundance). Potential sulfur-oxidizing bacteria were present at low relative abundances, including Sedimenticolaceae (bin_120, 0.2% and bin_221 HyVt-443, 0.1% rel. abundance) and Desulfobulbaceae (bin_064, 0.5% rel. abundance).

### Diverse and unique aerobic methanotrophs on carbonate rocks

16S rRNA sequence analysis of carbonate surfaces identified distinct aerobic methanotrophic lineages (Fig. 7). This included diverse members of Methylococcales clades that remained unclassified based on a phylogenetic tree (Fig. S9), that we refer to as uncultivated Methylomonadaceae-1, −2, −3 and −4. The Methylomonadaceae-3 subclade was recovered from several rock surfaces, including the SMM800 carbonates. IheB2-23 members were recovered from the R1 surface and, at high relative abundance, from a hydroid microbiome, attached to Del Mar R7 (Fig. 7d). IheB2-23 has previously been found on crabs [62], suggesting IheB2-23 may associate with seep invertebrates. Similarly, members of the Marine Methylotrophic Group 2 were recovered from all Del Mar carbonate surfaces, a lineage forming symbioses with sponges and seep-associated feather duster worms [63, 64]. Other aerobic methanotrophs appeared to have different habitat preferences. *Methyloprofundus*, a genus isolated from deep-sea sediments [65], was the main aerobic methanotroph recovered from Del Mar surface sediment, and from mud-like material in carbonates R3 and R4 (Fig. 7b,c), but had low relative abundance on carbonate surfaces. In the water column we mainly found OPU-1 and OPU-3 with a higher OPU-3 ratio at SMM800 associated with lower oxygen concentrations (Fig. 7a), consistent with earlier reports from oxygen minimum zones [66].

**Figure 7.**
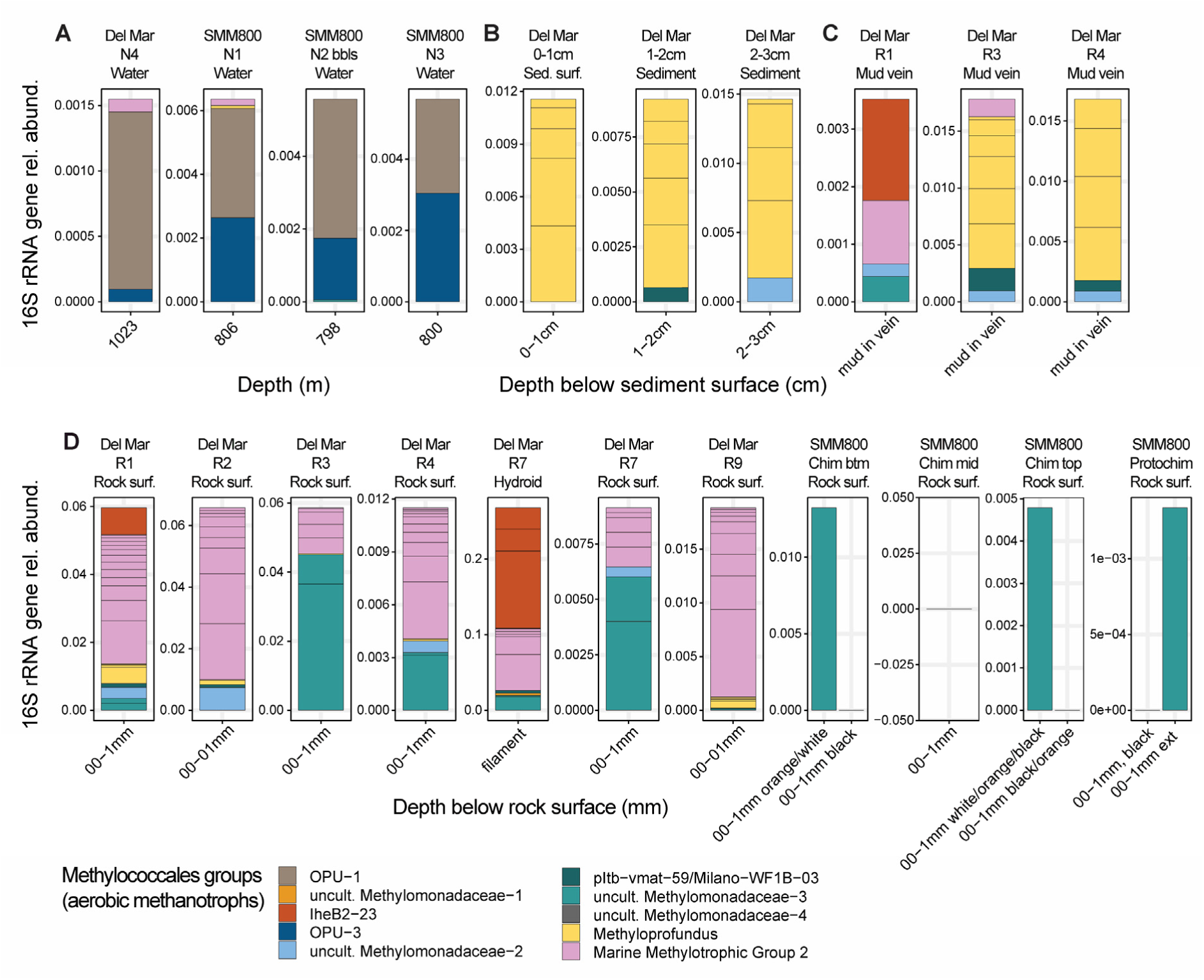
Aerobic methane-oxidizing bacteria on the carbonate surface compared to other seep habitats based on 16S rRNA gene sequencing. The carbonate surface harbors a distinct aerobic methanotrophic community. 16S rRNA sequence of the water column (A), the sediment surface (B), the mud in open vein (C), and the rock surface (D). We first classified the 16S rRNA sequences with SILVA and selected Methylococcales. Classification was refined with a 16S rRNA gene tree and reference sequences (Fig. S9).

### Carbonate-associated Methylophagaceae and unbinned contigs encode novel CuMMOs

Metagenomic analysis revealed an unusual diversity of carbonate-associated copper membrane monooxygenase genes (CuMMOs). CuMMOs are encoded by *xmoCAB* and catalyze aerobic oxidation of methane (*pmoCAB*), short-chain alkanes and ammonia [67]. Surprisingly, a member of Methylophagaceae (bin_133, genus GCA-002733105) encoded *xmoCAB*, to our knowledge the first reported CuMMO within Methylophagaceae (Fig. 8). Aerobic methanotrophs oxidize methane to methanol using *pmoCAB*, and Methylophagaceae are known methanol oxidizers. Therefore, methane oxidation with *xmoCAB* in Methylophagaceae appears likely. This is challenged by the finding that its *xmoC* is divergent from aerobic methanotrophs (Fig. 8), and oxidation of short-chain alkanes, ammonia or other substrates are equally likely hypothetical functions. Further *xmoC* diversity was recovered from R1 surface assemblies and co-assemblies (unbinned contigs). A new *xmoC* clade clustered with *xmoC* from *Nocardioides* CF8 and *Mycolicibacterium chubuense* (Actinomycetes), both known short-chain alkane (propane/butane) oxidizers [68, 69]. Two sequences clustered with ETHIRO and *Cycloclasticus* (Proteobacteria), that likely oxidize ethane or propane [70], and one *xmoC* clustered with *Halioglobus*. We found one *pxmC* sequence, a divergent *pmoC* with unknown function, commonly found in aerobic methanotrophs [71]. Additional to divergent *xmoC*s, we found conventional *pmoCs* (>40) clustering with methane-oxidizing Methylomonadaceae (Fig. 8, uncollapsed tree Fig. S10), and a clade clustering with but different from USCg, an aerobic methanotroph outside the order Methylococcales.

**Figure 8.**
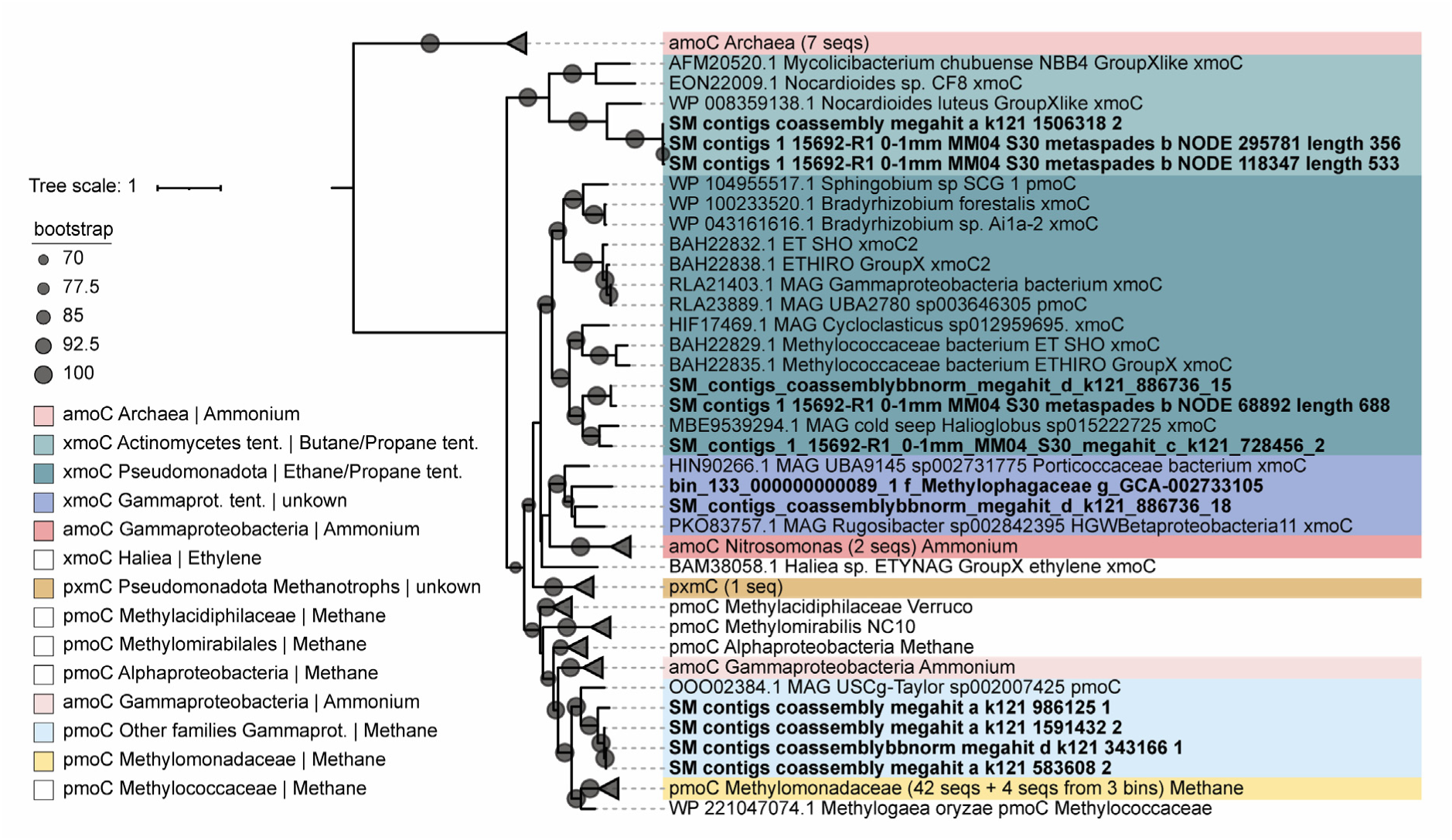
Phylogenetic tree of CuMMO subunit C gene (xmoC) recovered from carbonate-associated MAGs and metagenomic assemblies, as well as reference sequences. The clades for ammonium (*amoC*) and methane CuMMO (*pmoC*) are well supported by experimental data. Although *Mycolicibacterium* (synonym *Mycobacterium*) has been shown to oxidize butane, and ETHIRO (unpublished, isolate lost) has been suggested to oxidize ethane, the diversity within these groups is large, and especially for the seep sequences the substrates represent hypotheses. The *xmoC* Gammaprot. tent. (darker blue) has an unknown substrate so far but represents the first CuMMO within Methylophagaceae. Pmo refers to the genes encoding particulate methane monooxygenase, pxm to a version that some aerobic methanotrophs encode with unknown function, amo to the ammonia monooxygenase, and xmo to genes encoding CuMMO more generally. Sequences that were recovered from individual assemblies or co-assemblies, and were not part of a MAG, could not be assigned to specific taxonomic groups. Note that Proteobacteria and Pseudomonadota refer to the same phylum. Collapsed clades are labeled with the number of sequences recovered in this study (see uncollapsed tree Fig. S10). Abbreviations: seqs, sequences; tent., tentatively.

## Discussion

Here we demonstrate methane seep carbonate rocks host distinct surface and interior microbial communities both with previously unrecognized ecophysiology and diversity, including *Ca*. Desulfaltia, a potential new clade of ANME partner bacteria, and a potential new metabolic capability within Methylophagaceae containing a CuMMO (Fig. 3, 8). We further show that the endolithic carbonate AOM community is resilient [72], maintaining viability and the capacity to serve as methane sink over periods with fluctuating methane seepage (Fig. 6). The combination of geochemical, cell-specific activity alongside DNA-based analyses further demonstrated dormant/dead ANME cells occurring alongside active methanotrophic archaea (Fig. 5), revealing a new ecophysiological aspect of these rock-hosted archaea. We discovered a carbonate-surface ANME-SRB biofilm, previously only reported from of the Black Sea, but here, facing low-oxygen seawater. The carbonate surfaces further featured a sulfide-oxidizing bacterial mat, and another community without dominant members, with implications for carbonate dissolution/precipitation. The surface further harbored a distinct aerobic methanotroph community, that represents a potential pool for animal epibionts (Fig. 7). Novel carbonate-associated CuMMOs suggest these enzymes are more diverse and encoded by a wider range of microbes than previously recognized.

### Ecological insights of carbonate-hosted ANME-1 and their potential SRB partners

ANME-1 members are abundant in many seep carbonates [16], but an investigation of endolithic ANME-1 genera had been lacking. The prevalent genus *Ca.* Methanophaga has been recovered from cold seep sediments and seep carbonates [18, 73, 74]. We recovered two additional genera, QENJ01 and QEXZ01, also found in cold seep and hydrothermal vent sediments [74, 75]. *Ca*. Methanoalium-2 (JACGMN01), detected in our 16S rRNA survey at low relative abundance, occurs in marine seeps and terrestrial sites [73, 74, 76]. *Ca*. Methanoalium MAGs contain hydrogenases unlike related ANME and *Ca*. Methanoalium has been proposed to be methanogenic [73].

Alongside ANME-1, *Ca*. Desulfaltia (Seep-SRB-1b) with unknown ecology [77], was the most enriched SRB inside carbonates (Fig. 3), which we hypothesize may represent a new ANME-1 syntrophic partner. This group is a sister lineage to syntrophic Seep-SRB-1a (Fig. 4), and MAGs recovered from methane seeps, euxinic Black Sea waters and groundwater [78–81], often co-occured with ANME. The active endolithic ANME-1 in our study were primarily recovered as single cells rather than aggregates, but it is difficult to rule out disaggregation of loosely associated consortia during cell separation. Follow-up studies are needed to confirm its syntrophic partnership with ANME-1, however, in our genome analysis of related *Ca*. Desulfaltia MAGs we observed encoded multiheme cytochromes (*oetAB*), predicted to be used in direct interspecies electron transfer with ANME [58].

### Activity and viability of ANME cells in carbonates with low AOM

This study covers a large range of carbonate-associated AOM rates. Chimlet (840 – 3200 nmol cm^−3^ d^−1^) had comparable rates to the highest carbonate AOM rates (440 – 5500 nmol cm^−3^ d^−1^) reported from Point Dume, another Southern California seep [14], while the Del Mar outcrop carbonates had rates comparable to the lowest rates reported in [14] (5 nmol cm^−3^ d^−1^). Unexpectedly, low AOM carbonates maintained high 16S and metagenome-based ANME relative abundances, and BONCAT-FISH assays suggested ANME cells were rare, dormant or dead. Additionally, extracellular DNA may explain the observed ANME-DNA in low AOM carbonates, potentially well preserved through the sheltered environment, buffered pH and low temperature [82, 83]. On the other hand, extracellular DNA cannot completely explain the detected ANME-DNA as the reactivation of low AOM activity carbonates (Fig. 6) suggested that few (because of the long lag time) remaining viable AOM microorganisms reactivated AOM activity via exponential growth.

### Carbonates are potential long-term methane sinks

Methane seepage fluctuates [9], and reactivation of low AOM seep carbonates with renewed methane exposure had not previously been shown. Here, we experimentally reactivated low AOM carbonates with methane, showing carbonates remain potential ANME-SRB habitats and methane sinks long-term. The apparent 44-day doubling time was similar to the fastest reported ANME-SRB doubling time of 0.7-1 months measured in sediment from a seep-periphery [84] (literature comparison, supplementary table 4). Presumably low initial ANME-SRB cell numbers in our low AOM carbonates and the seep periphery sediment may allow fast growth through relaxed resource or space limitations. We further speculate that the present-day carbonate-hosted community is not the primary ANME-SRB community that once precipitated the carbonate. Given that carbonates precipitate over hundreds to thousands of years with layers of different ages [3], a successively changing ANME-SRB community may have facilitated precipitation.

### Metabolic range and role of the carbonate surface community

Anaerobes typically do not tolerate prolonged oxygen exposure [85]. Hence, the black-colored ANME-SRB surface-biofilms on the SMM800 carbonates occurring in direct contact with the overlying oxic seawater was unexpected (Fig. 2). Site SMM800 has high methane seepage and occurs within an oxygen minimum zone (8 µM oxygen), which may explain their ability to colonize the chemoherm surfaces. Co-occurring sulfide-oxidizing bacteria may further promote anoxic conditions at the surface [19, 86]. These biofilms resemble the microbial reefs described from the euxinic Black Sea seeps with cm-scale ANME-dominated mats [87]. Recently, ANME and related methanogens were hypothesized to produce black amorphous carbon [88]. Here, the black color of the ANME-SRB biofilm dissipated over time in fixative, arguing against amorphous carbon, and for unstable pigments or iron-sulfide minerals.

Sulfide-oxidizers and aerobic methanotrophs produce acidity that can dissolve seep carbonates [20, 21]. Our in-depth characterization of surface-associated microbial communities across different seep activities suggests the metabolic potential for dissolution is variable. Specifically, communities with less sulfide-oxidizing bacteria that were associated with low AOM carbonates, as described from carbonate colonization experiments [16], likely have a lower carbonate dissolution potential than sulfide-oxidizer dominated communities like on R9 (Fig. 1, 3). The ANME-SRB biofilms at SMM800 may continue to precipitate carbonate and protect the carbonate surface from corrosive co-occurring sulfide-oxidizers. Targeted investigations are needed to better constrain the spatial extent, environmental context, and contribution of surface microbial communities to carbonate dissolution and precipitation in the deep sea.

Microbial lineages were found to overlap between carbonate and animal surfaces. For example, the Del Mar carbonates shared the aerobic methanotrophic lineages IheB2-23 and MMG-2 and the methylotrophic group Methylophagaceae with carbonate-associated seep invertebrates, including hydroids (Fig. 7; S9) and sea spiders (pycnogonads) [62, 89, 90]. These shared taxa point to the potential importance of the carbonate surface community in animal epibiont recruitment and exchange.

Here we recovered a Methylophagacea MAG of the genus GCA-002733105 (bin_133) encoding a CuMMO with homology to methane, ammonia and hydrocarbon CuMMOs (Fig. 8). GCA-002733105 without CuMMO have been described, e.g., as symbionts within bathymodioline mussels supported by C1-compounds from methanotrophs [91]. We speculate GCA-002733105 members encoding CuMMO may have a methane-oxidizing potential, as methylotrophic capabilities are present and well documented within Methylophagaceae [92, 93]. The Methylophagaceae *xmoC* falls outside known methanotrophic *pmoC*s, suggesting a divergent evolutionary history. Alternatively, e.g., ammonia or short-chain alkanes may be likely alternative substrates. Future cultivation or enrichment are needed to determine taxonomy and substrate specificity of the recovered divergent *xmoCs* (Fig. 8), which would yield a more complete picture of CuMMO evolution and function.

In this study, we advanced the understanding of the diversity of microorganisms, their metabolic potential and activities within and on the surface of seep carbonates over a range of AOM activities. We identified a potential new ANME-partner *Ca*. Desulfaltia, which together with further validation, might expand the known diversity of ANME-SRB partnerships. Our results emphasize that DNA sequences do not always equate microbial activity, and ecophysiological measurements allow deeper insights into dynamics and physiological states of environmental microbes. By reactivating low AOM carbonates, we showed that seep carbonates remain potential ANME-SRB habitats, even over ceasing and recurring methane seepage, acting as potential methane sinks over carbonate lifetimes of 1000s of years. Further, we revealed the carbonate surface community as a distinct seep assemblage that deserves further attention, with potential to play a role in carbonate precipitation or dissolution, as a possible reservoir for animal epibionts, and as a host for novel CuMMO diversity. Finally, carbonate-hosted abundant and active microbes raise the question if and which other rock types in the deep sea and beyond may host microbial communities contributing to the global elemental cycles.

## Data availability

All raw reads generated in this study and selected high quality MAGs are available from NCBI under project number PRJNA1196099. All MAGs are available from FigShare.

## Acknowledgements

We are grateful to the R/V Western Flyer crew (Monterey Bay Research Aquarium Institute), John Magyar, Rebecca Wipfler, Sujung Lim and Shana Goffredi for sample retrieval. We thank Kriti Sharma for advice on rock work and BONCAT, Dan Utter for advice on bioinformatics, and Lydia Varesio and the ecology reading group for their thoughtful comments on this manuscript. We acknowledge Makayla Betts, Alex Sessions and the Resnick Sustainability Institute’s Water and Environment Lab at Caltech for isotope measurement support. This research was supported by: NSF project (2048666) to V.J.O. and SNF Postdoc Fellowship (P2EZP3_195375) to M.J.M.

## Author contributions

MJM and VJO designed the study and acquired funding. MJM was responsible for conducting the research, carried out laboratory experiments, data analyses and wrote the original manuscript. SAP contributed to rock preparation and sediment 16S rRNA sequencing. SAC performed amplicon libraries and FISH probe design. AKN performed the metagenomics libraries. RM contributed initial trees, sequence databases and data interpretation. AC contributed to data interpretation and guided biogeochemical analyses. VJO supervised the research and provided the samples and research infrastructure. All authors contributed to writing, reviewing and editing the manuscript.

## References

1. Knittel K, Boetius A. Anaerobic Oxidation of Methane: Progress with an Unknown Process. Annual Review of Microbiology 2009; 63: 311–334; doi: 10.1146/annurev.micro.61.080706.093130.

2. Blouet J-P, Arndt S, Imbert P, Regnier P. Are seep carbonates quantitative proxies of CH4 leakage? Modeling the influence of sulfate reduction and anaerobic oxidation of methane on pH and carbonate precipitation. Chemical Geology 2021; 577: 120254; doi: 10.1016/j.chemgeo.2021.120254.

3. Liebetrau V, Augustin N, Kutterolf S, Schmidt M, Eisenhauer A, Garbe-Schönberg D, et al. Cold-seep-driven carbonate deposits at the Central American forearc: contrasting evolution and timing in escarpment and mound settings. Int J Earth Sci (Geol Rundsch) 2014; 103: 1845–1872; doi: 10.1007/s00531-014-1045-2.

4. Crémière A, Lepland A, Chand S, Sahy D, Kirsimäe K, Bau M, et al. Fluid source and methane-related diagenetic processes recorded in cold seep carbonates from the Alvheim channel, central North Sea. Chemical Geology 2016; 432: 16–33; doi: 10.1016/j.chemgeo.2016.03.019.

5. Schroedl P, Silverstein M, DiGregorio D, Blättler CL, Loyd S, Bradbury HJ, et al. Carbonate chimneys at the highly productive point Dume methane seep: Fine-scale mineralogical, geochemical, and microbiological heterogeneity reflects dynamic and long-lived methane-metabolizing habitats. Geobiology 2024; 22: e12608; doi: 10.1111/gbi.12608.

6. Akam SA, Swanner ED, Yao H, Hong W-L, Peckmann J. Methane-derived authigenic carbonates – A case for a globally relevant marine carbonate factory. Earth-Science Reviews 2023; 243: 104487; doi: 10.1016/j.earscirev.2023.104487.

7. Danovaro R, Levin LA, Fanelli G, Scenna L, Corinaldesi C. Microbes as marine habitat formers and ecosystem engineers. Nat Ecol Evol 2024; 8: 1407–1419; doi: 10.1038/s41559-024-02407-7.

8. Levin LA, Mendoza GF, Grupe BM. Methane seepage effects on biodiversity and biological traits of macrofauna inhabiting authigenic carbonates. Deep Sea Research Part II: Topical Studies in Oceanography 2017; 137: 26–41; doi: 10.1016/j.dsr2.2016.05.021.

9. Tryon MD, Brown KM, Torres ME. Fluid and chemical flux in and out of sediments hosting methane hydrate deposits on Hydrate Ridge, OR, II: Hydrological processes. Earth and Planetary Science Letters 2002; 201: 541–557; doi: 10.1016/S0012-821X(02)00732-X.

10. Thauer RK, Shima S. Methane as Fuel for Anaerobic Microorganisms. Annals of the New York Academy of Sciences 2008; 1125: 158–170; doi: 10.1196/annals.1419.000.

11. Orphan VJ, Turk KA, Green AM, House CH. Patterns of 15N assimilation and growth of methanotrophic ANME-2 archaea and sulfate-reducing bacteria within structured syntrophic consortia revealed by FISH-SIMS. Environmental Microbiology 2009; 11: 1777–1791; doi: 10.1111/j.1462-2920.2009.01903.x.

12. Guan H, Birgel D, Peckmann J, Liang Q, Feng D, Yang S, et al. Lipid biomarker patterns of authigenic carbonates reveal fluid composition and seepage intensity at Haima cold seeps, South China Sea. Journal of Asian Earth Sciences 2018; 168: 163–172; doi: 10.1016/j.jseaes.2018.04.035.

13. Wang Y, Li W, Li Q, Zhou Y, Gao Z, Feng D. Deep-Sea Carbonates Are a Reservoir of Fossil Microbes Previously Inhabiting Cold Seeps. Frontiers in Marine Science 2021; 8: 698945; doi: 10.3389/fmars.2021.698945.

14. Marlow JJ, Hoer D, Jungbluth SP, Reynard LM, Gartman A, Chavez MS, et al. Carbonate-hosted microbial communities are prolific and pervasive methane oxidizers at geologically diverse marine methane seep sites. PNAS 2021; 118; doi: 10.1073/pnas.2006857118.

15. Marlow JJ, Steele JA, Ziebis W, Thurber AR, Levin LA, Orphan VJ. Carbonate-hosted methanotrophy represents an unrecognized methane sink in the deep sea. Nature Communications 2014; 5: 5094; doi: 10.1038/ncomms6094.

16. Case DH, Pasulka AL, Marlow JJ, Grupe BM, Levin LA, Orphan VJ. Methane seep carbonates host distinct, diverse, and dynamic microbial assemblages. MBio 2015; 6.

17. Lee D-H, Kim J-H, Lee YM, Bayon G, Kim D, Joe YJ, et al. Metalloenzyme signatures in authigenic carbonates from the Chukchi Borderlands in the western Arctic Ocean. Sci Rep 2022; 12: 16597; doi: 10.1038/s41598-022-21184-6.

18. Beckmann S, Farag IF, Zhao R, Christman GD, Prouty NG, Biddle JF. Expanding the repertoire of electron acceptors for the anaerobic oxidation of methane in carbonates in the Atlantic and Pacific Ocean. The ISME Journal 2021; 1–14; doi: 10.1038/s41396-021-00918-w.

19. Bailey JV, Orphan VJ, Joye SB, Corsetti FA. Chemotrophic microbial mats and their potential for preservation in the rock record. Astrobiology 2009; 9: 843–859; doi: 10.1089/ast.2008.0314.

20. Leprich DJ, Flood BE, Schroedl PR, Ricci E, Marlow JJ, Girguis PR, et al. Sulfur bacteria promote dissolution of authigenic carbonates at marine methane seeps. The ISME Journal 2021; 1–14; doi: 10.1038/s41396-021-00903-3.

21. Cordova-Gonzalez A, Birgel D, Wisshak M, Urich T, Brinkmann F, Marcon Y, et al. A carbonate corrosion experiment at a marine methane seep: The role of aerobic methanotrophic bacteria. Geobiology 2023; 21: 491–506; doi: 10.1111/gbi.12549.

22. Levin LA, Mendoza GF, Grupe BM, Gonzalez JP, Jellison B, Rouse G, et al. Biodiversity on the Rocks: Macrofauna Inhabiting Authigenic Carbonate at Costa Rica Methane Seeps. PLoS ONE 2015; 10: e0131080; doi: 10.1371/journal.pone.0131080.

23. Grupe BM, Krach ML, Pasulka AL, Maloney JM, Levin LA, Frieder CA. Methane seep ecosystem functions and services from a recently discovered southern California seep. Marine Ecology 2015; 36: 91–108; doi: 10.1111/maec.12243.

24. Paull CK, Normark WR, Ussler W, Caress DW, Keaten R. Association among active seafloor deformation, mound formation, and gas hydrate growth and accumulation within the seafloor of the Santa Monica Basin, offshore California. Marine Geology 2008; 250: 258–275; doi: 10.1016/j.margeo.2008.01.011.

25. Marlow JJ, Steele JA, Ziebis W, Scheller S, Case D, Reynard LM, et al. Monodeuterated methane, an isotopic tool to assess biological methane metabolism rates. mSphere 2017; 2; doi: 10.1128/mSphereDirect.00309-17.

26. Scheller S, Yu H, Chadwick GL, McGlynn SE, Orphan VJ. Artificial electron acceptors decouple archaeal methane oxidation from sulfate reduction. Science 2016; 351: 703–707; doi: 10.1126/science.aad7154.

27. Mullin SW, Wanger G, Kruger BR, Sackett JD, Hamilton-Brehm SD, Bhartia R, et al. Patterns of in situ mineral colonization by microorganisms in a ∼60°C deep continental subsurface aquifer. Front Microbiol 2020; 11; doi: 10.3389/fmicb.2020.536535.

28. Parada AE, Needham DM, Fuhrman JA. Every base matters: assessing small subunit rRNA primers for marine microbiomes with mock communities, time series and global field samples. Environmental Microbiology 2016; 18: 1403–1414; doi: 10.1111/1462-2920.13023.

29. Seemann T. barrnap https://github.com/tseemann/barrnap.

30. Minh BQ, Schmidt HA, Chernomor O, Schrempf D, Woodhams MD, von Haeseler A, et al. IQ-TREE 2: New models and efficient methods for phylogenetic inference in the genomic era. Molecular Biology and Evolution 2020; 37: 1530–1534; doi: 10.1093/molbev/msaa015.

31. Bushnell B. BBMap sourceforge.net/projects/bbmap/.

32. Li D, Liu C-M, Luo R, Sadakane K, Lam T-W. MEGAHIT: an ultra-fast single-node solution for large and complex metagenomics assembly via succinct de Bruijn graph. Bioinformatics 2015; 31: 1674–1676; doi: 10.1093/bioinformatics/btv033.

33. Nurk S, Meleshko D, Korobeynikov A, Pevzner PA. metaSPAdes: a new versatile metagenomic assembler. Genome Res 2017; 27: 824–834; doi: 10.1101/gr.213959.116.

34. Uritskiy GV, DiRuggiero J, Taylor J. MetaWRAP—a flexible pipeline for genome-resolved metagenomic data analysis. Microbiome 2018; 6: 158; doi: 10.1186/s40168-018-0541-1.

35. Olm MR, Brown CT, Brooks B, Banfield JF. dRep: a tool for fast and accurate genomic comparisons that enables improved genome recovery from metagenomes through de-replication. The ISME Journal 2017; 11: 2864–2868; doi: 10.1038/ismej.2017.126.

36. Eren AM, Esen ÖC, Quince C, Vineis JH, Morrison HG, Sogin ML, et al. Anvi’o: an advanced analysis and visualization platform for ‘omics data. PeerJ 2015; 3: e1319; doi: 10.7717/peerj.1319.

37. Vollmers J, Wiegand S, Lenk F, Kaster A-K. How clear is our current view on microbial dark matter? (Re-)assessing public MAG & SAG datasets with MDMcleaner. Nucleic Acids Research 2022; 50: e76; doi: 10.1093/nar/gkac294.

38. Chaumeil P-A, Mussig AJ, Hugenholtz P, Parks DH. GTDB-Tk v2: memory friendly classification with the Genome Taxonomy Database. 2022. bioRxiv., 2022.07.11.499641

39. Woodcroft B. J. coverM https://github.com/wwood/CoverM).

40. Zhou Z, Tran PQ, Breister AM, Liu Y, Kieft K, Cowley ES, et al. METABOLIC: high-throughput profiling of microbial genomes for functional traits, metabolism, biogeochemistry, and community-scale functional networks. Microbiome 2022; 10: 33; doi: 10.1186/s40168-021-01213-8.

41. Parks DH, Imelfort M, Skennerton CT, Hugenholtz P, Tyson GW. CheckM: assessing the quality of microbial genomes recovered from isolates, single cells, and metagenomes. Genome Res 2015; 25: 1043– 1055; doi: 10.1101/gr.186072.114.

42. Chklovski A, Parks DH, Woodcroft BJ, Tyson GW. CheckM2: a rapid, scalable and accurate tool for assessing microbial genome quality using machine learning. Nat Methods 2023; 20: 1203–1212; doi: 10.1038/s41592-023-01940-w.

43. Laso-Pérez R, Wu F, Crémière A, Speth DR, Magyar JS, Zhao K, et al. Evolutionary diversification of methanotrophic ANME-1 archaea and their expansive virome. Nat Microbiol 2023; 1–15; doi: 10.1038/s41564-022-01297-4.

44. Hyatt D, Chen G-L, LoCascio PF, Land ML, Larimer FW, Hauser LJ. Prodigal: prokaryotic gene recognition and translation initiation site identification. BMC Bioinformatics 2010; 11: 119; doi: 10.1186/1471-2105-11-119.

45. Buchfink B, Xie C, Huson DH. Fast and sensitive protein alignment using DIAMOND. Nat Methods 2015; 12: 59–60; doi: 10.1038/nmeth.3176.

46. Sayers EW, Bolton EE, Brister JR, Canese K, Chan J, Comeau DC, et al. Database resources of the national center for biotechnology information. Nucleic Acids Res 2022; 50: D20–D26; doi: 10.1093/nar/gkab1112.

47. Letunic I, Bork P. Interactive Tree of Life (iTOL) v6: recent updates to the phylogenetic tree display and annotation tool. Nucleic Acids Research 2024; gkae268; doi: 10.1093/nar/gkae268.

48. Eichorst SA, Strasser F, Woyke T, Schintlmeister A, Wagner M, Woebken D. Advancements in the application of NanoSIMS and Raman microspectroscopy to investigate the activity of microbial cells in soils. FEMS Microbiology Ecology 2015; 91: fiv106; doi: 10.1093/femsec/fiv106.

49. Orphan VJ, House CH, Hinrichs K-U, McKeegan KD, DeLong EF. Multiple archaeal groups mediate methane oxidation in anoxic cold seep sediments. PNAS 2002; 99: 7663–7668; doi: 10.1073/pnas.072210299.

50. Mayr MJ, Orphan VJ. Extraction of microbial cells from methane-seep carbonate rocks for single-cell analyses (e.g. BONCAT-FISH) using “combination buffer” and percoll density centrifugation. 10.17504/protocols.io.rm7vzjk55lx1/v1.

51. Hatzenpichler R, Connon SA, Goudeau D, Malmstrom RR, Woyke T, Orphan VJ. Visualizing in situ translational activity for identifying and sorting slow-growing archaeal−bacterial consortia. PNAS 2016; 113: E4069–E4078; doi: 10.1073/pnas.1603757113.

52. Daims H, Brühl A, Amann R, Schleifer K-H, Wagner M. The domain-specific probe EUB338 is insufficient for the detection of all bacteria: Development and evaluation of a more comprehensive probe set. Systematic and Applied Microbiology 1999; 22: 434–444; doi: 10.1016/S0723-2020(99)80053-8.

53. Stahl, D. A., Amann, R. I. Development and application of nucleic acid probes in bacterial systematics. Nucleic Acid Techniques in Bacterial Systematics. 1991. Wiley.

54. Boetius A, Ravenschlag K, Schubert CJ, Rickert D, Widdel F, Gieseke A, et al. A marine microbial consortium apparently mediating anaerobic oxidation of methane. Nature 2000; 407: 623–626; doi: 10.1038/35036572.

55. Google. Map of Southern California. 2024. Google Maps.

56. Michaelis W, Seifert R, Nauhaus K, Treude T, Thiel V, Blumenberg M, et al. Microbial reefs in the Black Sea fueled by anaerobic oxidation of methane. Science 2002; 297: 1013–1015; doi: 10.1126/science.1072502.

57. Schreiber L, Holler T, Knittel K, Meyerdierks A, Amann R. Identification of the dominant sulfate-reducing bacterial partner of anaerobic methanotrophs of the ANME-2 clade. Environmental microbiology 2010; 12: 2327–2340; doi: 10.1111/j.1462-2920.2010.02275.x.

58. Murali R, Yu H, Speth DR, Wu F, Metcalfe KS, Crémière A, et al. Physiological potential and evolutionary trajectories of syntrophic sulfate-reducing bacterial partners of anaerobic methanotrophic archaea. PLOS Biology 2023; 21: e3002292; doi: 10.1371/journal.pbio.3002292.

59. Wegener G, Krukenberg V, Riedel D, Tegetmeyer HE, Boetius A. Intercellular wiring enables electron transfer between methanotrophic archaea and bacteria. Nature 2015; 526: 587–590; doi: 10.1038/nature15733.

60. Holler T, Widdel F, Knittel K, Amann R, Kellermann MY, Hinrichs K-U, et al. Thermophilic anaerobic oxidation of methane by marine microbial consortia. ISME J 2011; 5: 1946–1956; doi: 10.1038/ismej.2011.77.

61. Salman-Carvalho V, Fadeev E, Joye SB, Teske A. How Clonal Is Clonal? Genome Plasticity across Multicellular Segments of a “Candidatus Marithrix sp.” Filament from Sulfidic, Briny Seafloor Sediments in the Gulf of Mexico. Frontiers in Microbiology 2016; 7.

62. Watsuji T, Nakagawa S, Tsuchida S, Toki T, Hirota A, Tsunogai U, et al. Diversity and function of epibiotic microbial communities on the Galatheid crab, Shinkaia crosnieri. Microb Environ 2010; 25: 288–294; doi: 10.1264/jsme2.ME10135.

63. Goffredi SK, Tilic E, Mullin SW, Dawson KS, Keller A, Lee RW, et al. Methanotrophic bacterial symbionts fuel dense populations of deep-sea feather duster worms (Sabellida, Annelida) and extend the spatial influence of methane seepage. Sci Adv 2020; 6: eaay8562; doi: 10.1126/sciadv.aay8562.

64. Petersen JM, Wentrup C, Verna C, Knittel K, Dubilier N. Origins and Evolutionary Flexibility of Chemosynthetic Symbionts From Deep-Sea Animals. The Biological Bulletin 2012; 223: 123–137; doi: 10.1086/BBLv223n1p123.

65. Tavormina PL, Hatzenpichler R, McGlynn S, Chadwick G, Dawson KS, Connon SA, et al. Methyloprofundus sedimenti gen. nov., sp. nov., an obligate methanotroph from ocean sediment belonging to the ‘deep sea-1’ clade of marine methanotrophs. International Journal of Systematic and Evolutionary Microbiology 2015; 65: 251–259; doi: 10.1099/ijs.0.062927-0.

66. Tavormina PL, Ussler III W, Steele JA, Connon SA, Klotz MG, Orphan VJ. Abundance and distribution of diverse membrane-bound monooxygenase (Cu-MMO) genes within the Costa Rica oxygen minimum zone. Environmental Microbiology Reports 2013; 5: 414–423; doi: 10.1111/1758-2229.12025.

67. Tucci FJ, Rosenzweig AC. Direct Methane Oxidation by Copper- and Iron-Dependent Methane Monooxygenases. Chem Rev 2024; 124: 1288–1320; doi: 10.1021/acs.chemrev.3c00727.

68. Sayavedra-Soto LA, Hamamura N, Liu C-W, Kimbrel JA, Chang JH, Arp DJ. The membrane-associated monooxygenase in the butane-oxidizing Gram-positive bacterium Nocardioides sp. strain CF8 is a novel member of the AMO/PMO family. Environmental Microbiology Reports 2011; 3: 390–396; doi: 10.1111/j.1758-2229.2010.00239.x.

69. Coleman NV, Le NB, Ly MA, Ogawa HE, McCarl V, Wilson NL, et al. Hydrocarbon monooxygenase in Mycobacterium: recombinant expression of a member of the ammonia monooxygenase superfamily. ISME J 2012; 6: 171–182; doi: 10.1038/ismej.2011.98.

70. Chakraborty R, Borglin SE, Dubinsky EA, Andersen GL, Hazen TC. Microbial Response to the MC-252 Oil and Corexit 9500 in the Gulf of Mexico. Front Microbiol 2012; 3; doi: 10.3389/fmicb.2012.00357.

71. Tavormina PL, Orphan VJ, Kalyuzhnaya MG, Jetten MSM, Klotz MG. A novel family of functional operons encoding methane/ammonia monooxygenase-related proteins in gammaproteobacterial methanotrophs. Environmental Microbiology Reports 2011; 3: 91–100; doi: 10.1111/j.1758-2229.2010.00192.x.

72. Shade A, Peter H, Allison SD, Baho D, Berga M, Buergmann H, et al. Fundamentals of Microbial Community Resistance and Resilience. Front Microbiol 2012; 3; doi: 10.3389/fmicb.2012.00417.

73. Chadwick GL, Skennerton CT, Laso-Pérez R, Leu AO, Speth DR, Yu H, et al. Comparative genomics reveals electron transfer and syntrophic mechanisms differentiating methanotrophic and methanogenic archaea. PLoS Biol 2022; 20: e3001508; doi: 10.1371/journal.pbio.3001508.

74. Vulcano F, Hahn CJ, Roerdink D, Dahle H, Reeves EP, Wegener G, et al. Phylogenetic and functional diverse ANME-1 thrive in Arctic hydrothermal vents. FEMS Microbiology Ecology 2022; 98: fiac117; doi: 10.1093/femsec/fiac117.

75. Hwang Y, Roux S, Coclet C, Krause SJE, Girguis PR. Viruses interact with hosts that span distantly related microbial domains in dense hydrothermal mats. Nat Microbiol 2023; 8: 946–957; doi: 10.1038/s41564-023-01347-5.

76. Parks DH, Chuvochina M, Rinke C, Mussig AJ, Chaumeil P-A, Hugenholtz P. GTDB: an ongoing census of bacterial and archaeal diversity through a phylogenetically consistent, rank normalized and complete genome-based taxonomy. Nucleic Acids Research 2022; 50: D785–D794; doi: 10.1093/nar/gkab776.

77. Kleindienst S, Knittel K. Anaerobic Hydrocarbon-Degrading Sulfate-Reducing Bacteria at Marine Gas and Oil Seeps. In: Teske A, Carvalho V (eds). Marine Hydrocarbon Seeps: Microbiology and Biogeochemistry of a Global Marine Habitat. 2020. Springer International Publishing, Cham, pp 21–41.

78. Ruff SE, Humez P, Angelis IH de, Nightingale M, Diao M, Cho S, et al. Hydrogen and dark oxygen drive microbial productivity in diverse groundwater ecosystems. 2022. bioRxiv., 2022.08.09.503387

79. Cabello-Yeves PJ, Callieri C, Picazo A, Mehrshad M, Haro-Moreno JM, Roda-Garcia JJ, et al. The microbiome of the Black Sea water column analyzed by shotgun and genome centric metagenomics. Environmental Microbiome 2021; 16: 5; doi: 10.1186/s40793-021-00374-1.

80. Dong X, Peng Y, Wang M, Woods L, Wu W, Wang Y, et al. Evolutionary ecology of microbial populations inhabiting deep sea sediments associated with cold seeps. Nat Commun 2023; 14: 1127; doi: 10.1038/s41467-023-36877-3.

81. Vigneron A, Alsop EB, Cruaud P, Philibert G, King B, Baksmaty L, et al. Comparative metagenomics of hydrocarbon and methane seeps of the Gulf of Mexico. Sci Rep 2017; 7: 16015; doi: 10.1038/s41598-017-16375-5.

82. Nielsen KM, Johnsen PJ, Bensasson D, Daffonchio D. Release and persistence of extracellular DNA in the environment. Environ Biosafety Res 2007; 6: 37–53; doi: 10.1051/ebr:2007031.

83. Wegner C-E, Stahl R, Velsko I, Hübner A, Fagernäs Z, Warinner C, et al. A glimpse of the paleome in endolithic microbial communities. Microbiome 2023; 11: 210; doi: 10.1186/s40168-023-01647-2.

84. Girguis PR, Cozen AE, DeLong EF. Growth and population dynamics of anaerobic methane-oxidizing archaea and sulfate-reducing bacteria in a continuous-flow bioreactor. Appl Environ Microbiol 2005; 71: 3725–3733; doi: 10.1128/AEM.71.7.3725-3733.2005.

85. Lu Z, Imlay JA. When anaerobes encounter oxygen: mechanisms of oxygen toxicity, tolerance and defence. Nat Rev Microbiol 2021; 19: 774–785; doi: 10.1038/s41579-021-00583-y.

86. Teichert BMA, Bohrmann G, Suess E. Chemoherms on Hydrate Ridge — Unique microbially-mediated carbonate build-ups growing into the water column. Palaeogeography, Palaeoclimatology, Palaeoecology 2005; 227: 67–85; doi: 10.1016/j.palaeo.2005.04.029.

87. Michaelis W, Seifert R, Nauhaus K, Treude T, Thiel V, Blumenberg M, et al. Microbial Reefs in the Black Sea Fueled by Anaerobic Oxidation of Methane. Science 2002; 297: 1013–1015; doi: 10.1126/science.1072502.

88. Allen KD, Wegener G, Matthew Sublett D, Bodnar RJ, Feng X, Wendt J, et al. Biogenic formation of amorphous carbon by anaerobic methanotrophs and select methanogens. Science Advances 2021; 7: eabg9739; doi: 10.1126/sciadv.abg9739.

89. Dal Bó, B., Guo, Y., Mayr, M. J., Pereira, O. S., Levin, L., Orphan, V. J., et al. Methane-powered sea spiders: Diverse, epibiotic methanotrophs serve as a novel source of nutrition for deep-sea methane seep Sericosura. under review.

90. Thurber AR, Jones WJ, Schnabel K. Dancing for food in the deep sea: bacterial farming by a new species of Yeti crab. PLOS ONE 2011; 6: e26243; doi: 10.1371/journal.pone.0026243.

91. Zvi-Kedem T, Vintila S, Kleiner M, Tchernov D, Rubin-Blum M. Metabolic handoffs between multiple symbionts may benefit the deep-sea bathymodioline mussels. ISME COMMUN 2023; 3: 1–13; doi: 10.1038/s43705-023-00254-4.

92. Boden R, Kelly DP, Murrell JC, Schäfer H. Oxidation of dimethylsulfide to tetrathionate by Methylophaga thiooxidans sp. nov.: a new link in the sulfur cycle. Environmental Microbiology 2010; 12: 2688–2699; doi: 10.1111/j.1462-2920.2010.02238.x.

93. de Zwart JMM, Nelisse PN, Kuenen JG. Isolation and characterization of Methylophaga sulfidovorans sp. nov.: an obligately methylotrophic, aerobic, dimethylsulfide oxidizing bacterium from a microbial mat. FEMS Microbiology Ecology 1996; 20: 261–270; doi: 10.1111/j.1574-6941.1996.tb00324.x.

