## Supplementary Figures for "Distinct Microbial Communities Within and On Seep Carbonates Support Long-term Anaerobic Oxidation of Methane and Novel pMMO Diversity"

**Supplementary Figure S1**

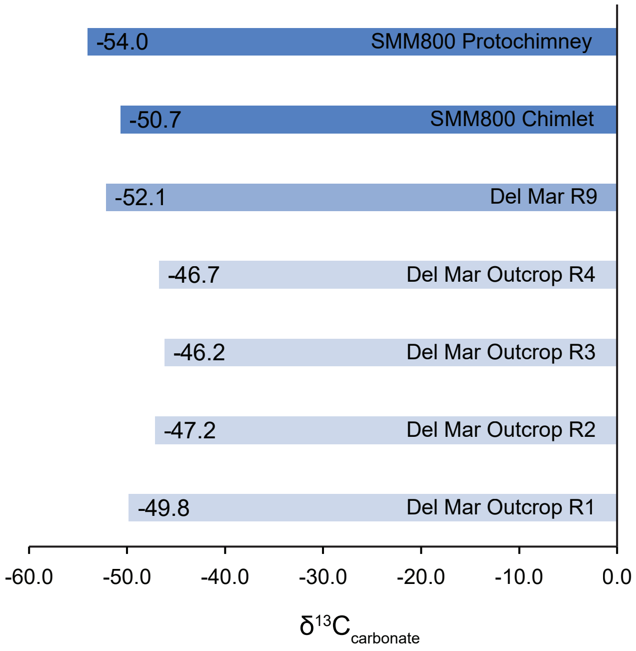


**Supplementary Figure S1 $\delta^{13}C_{carbonate}$ values of carbonate rocks from Del Mar and Santa Monica Mound 800 seeps.** The strongly depleted $\delta^{13}C_{carbonate}$ values measured from powdered carbonate clearly indicate a methane-derived origin of the carbonate rocks. The values are comparable between rocks with potentially slightly more depletion in the intermediate and high anaerobic methane oxidation activity rocks R9 as well as chimlet and protochimney.

### Supplementary Figure S2

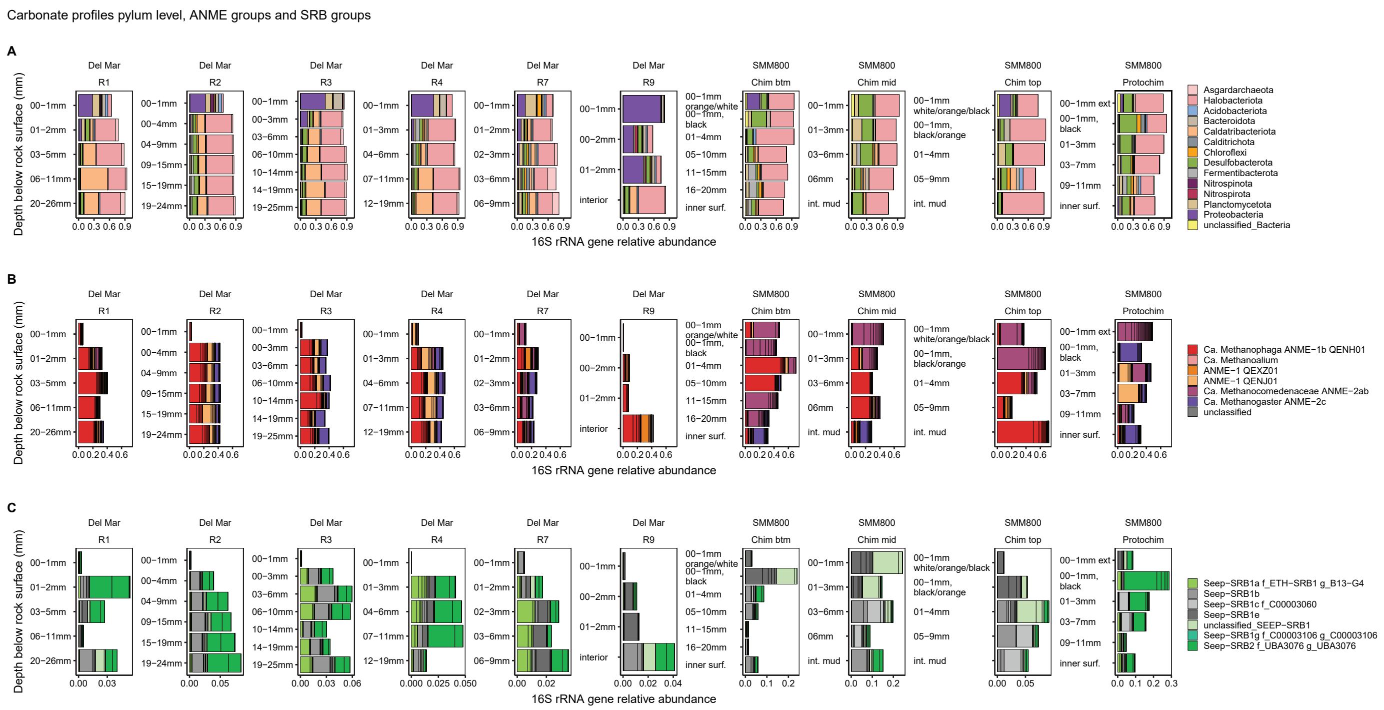

**Supplementary Figure S2 Microbial community from carbonate surface to interior in the seep context.** 16S rRNA gene sequencing of carbonate surface scrapes and mm-scale sections. **A**) Archaeal and bacterial phyla in selected sectioned carbonates from surface to interior based on 16S rRNA gene sequencing. Phyla reaching ≥5% at least once are shown. Carbonates naturally varied in size and shape which led to different section sizes, even though we tried to cut them as similarly as possible. We sectioned three parts of chimlet: bottom (btm), middle (mid), top (cap) **B**) ANME phylogenetic groups based on gtdb taxonomy identified with a 16S rRNA phylogenetic tree to genus level where possible. **C**) SRB phylogenetic groups based a 16S rRNA phylogenetic tree. For ANME and SRB ASVs reaching ≥1% at least once were included. Note that SRB x-axes are scaled to the maximum abundance of the respective profile for better visibility, because SRB abundances varied substantially. Abbreviations: Inner surf., inner surface of chimlet and protochimney cavity; int, interior;

**Supplementary Figure S3**

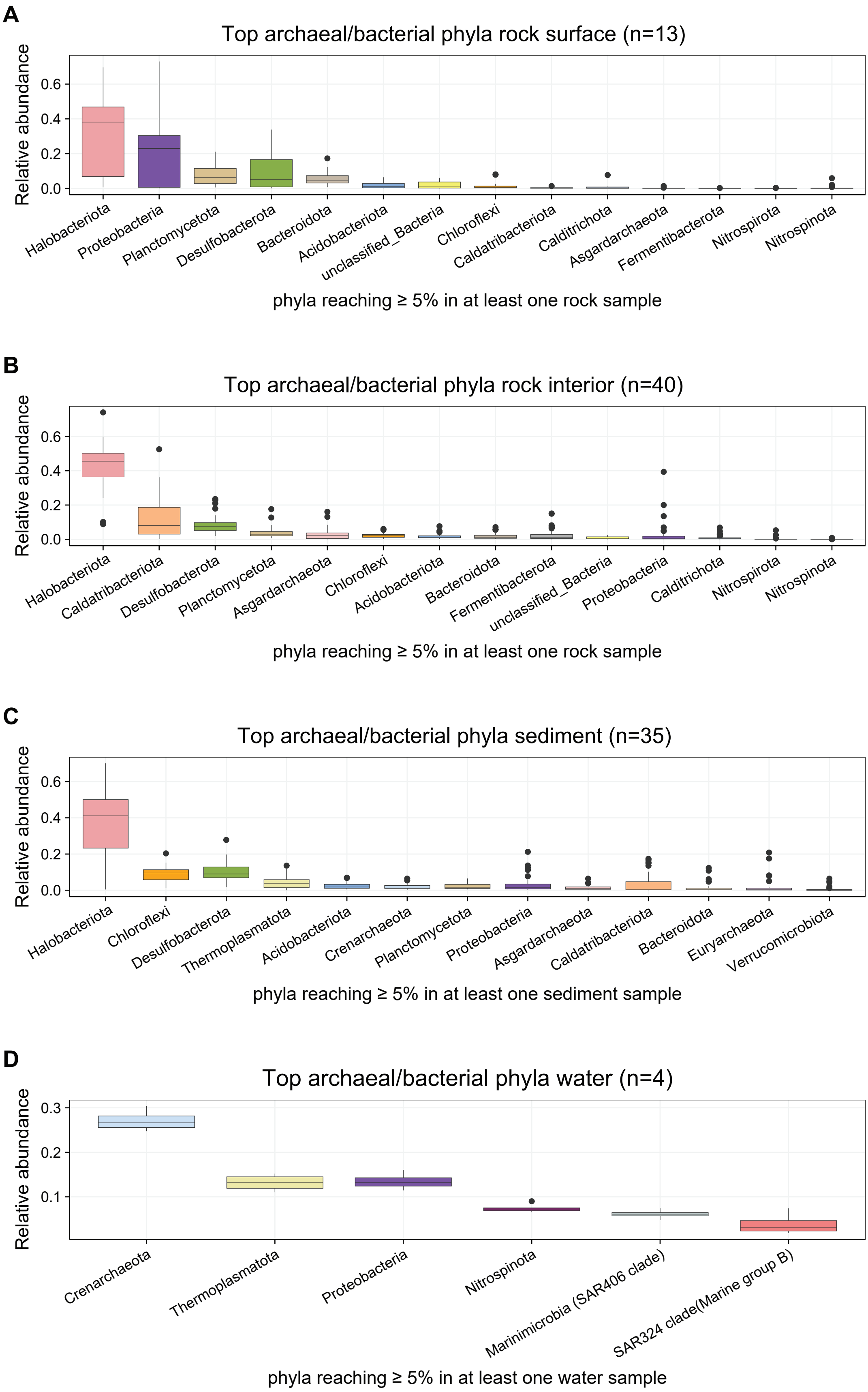


**Supplementary Figure S3 Top bacterial and archaeal phyla in different seep environments.** Phyla reaching ≥5% at least once are shown. The top most sediment layer was not included in this analysis. Note that the y-axis of the water column has a different scale for better visibility. The number of samples (n) that were included to calculate the median is shown in figure headers. **A**) rock surface **B**) rock interior **C**) sediment and **D**) water column.

### Supplementary Figure S4

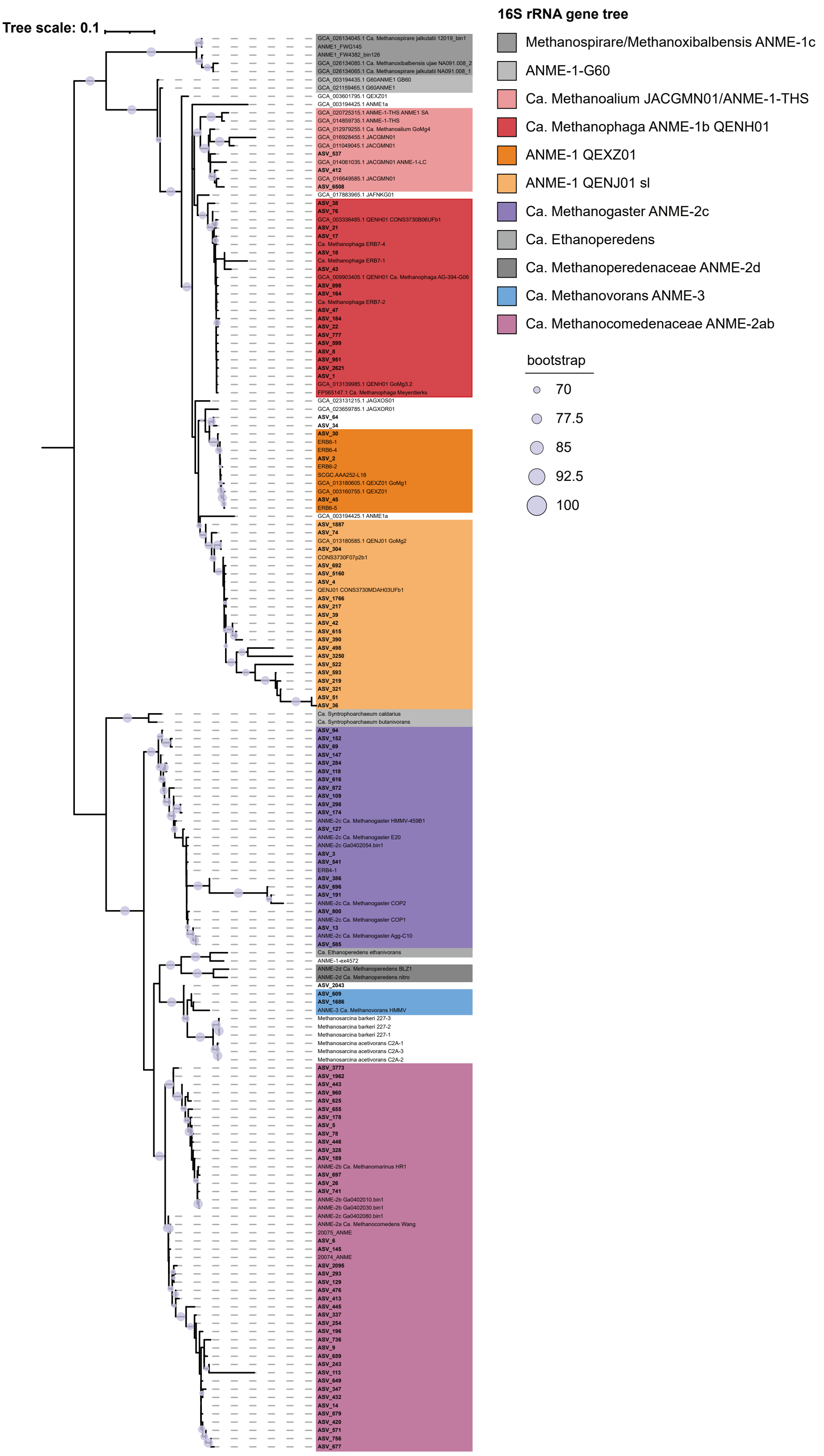


**Supplementary Figure S4 16S rRNA phylogenetic tree of ANME ASVs found in this study with reference sequences from gtdb to further classify ANME-1 (Methanospirareceae) beyond SILVA ANME-1a and ANME-1b.** 16S rRNA sequences were first classified with SILVA. 16S rRNA sequences from gtdb were added starting with GCA. Sequences from this study in bold. In case of QEXZ01 one representative does not cluster with the rest of the sequences (GCA_003601795.1 QEXZ01), because GoMg1 is a SAG (single amplified genome) we trusted its 16S rRNA sequence. Binning 16S rRNA genes correctly can be difficult, therefore some noise in 16S rRNA genes binned with MAGs can be expected. ANME-3 were not found in carbonate rocks above 1%, the ANME-3 ASVs shown in the tree were above 1% in some sediment samples. A few additional inhouse 16S rRNA sequences were added for our reference (names without GCA). For ANME-2c we used the SILVA classification. For ANME-2ab we stopped at family level, because the SILVA genus grouping into ANME-2ab and ANME-2b is ambiguous and not clearly resolved in the tree here.

### Supplementary Figure S5

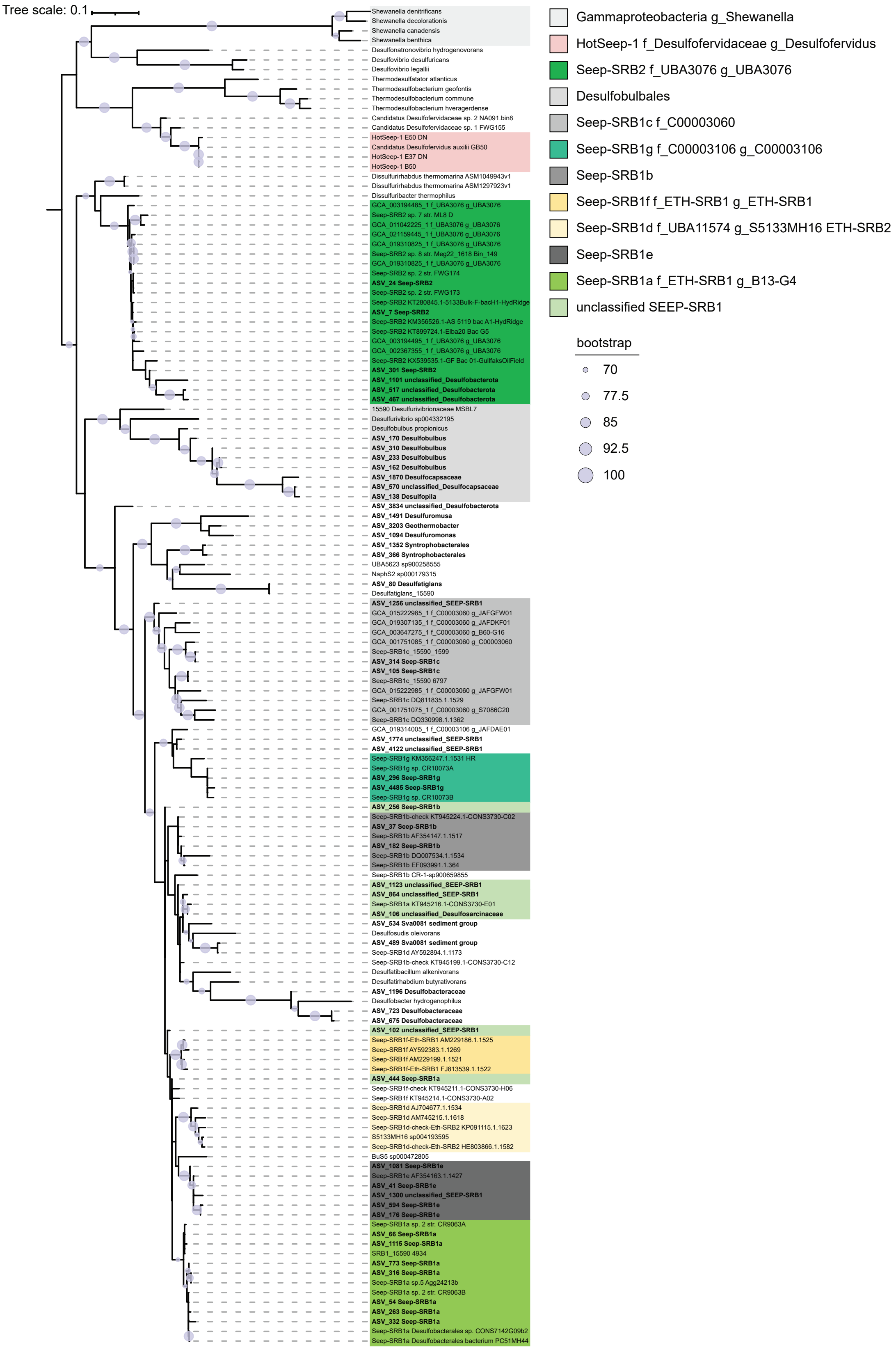


**Supplementary Figure S5 16S rRNA phylogenetic tree of Seep-SRB1 and 2 ASVs found in this study with reference sequences.** Seep-SRB1a, Seep-SRB1g and Seep-SRB2 are currently known SRB partner bacteria of ANME. HotSeep-1 is another group of partner bacteria not found in this study. 16S rRNA sequences were first classified with SILVA and an inhouse SRB database (https://doi.org/10.6084/m9.figshare.27288090.v1). We checked the classification of Desulfobacterota ASVs,reaching ≥1% in at least one sample with this tree and updated a few classifications according to the tree. The names next to the sequence is the original classification, the colors depict the slightly changed manual classification based on the tree. The tree is not resolving all groups well, and we tried to classify cautiously. The legend shows the gtdb taxonomy where available. ASVs found in this study in bold.

**Supplementary Figure S6**

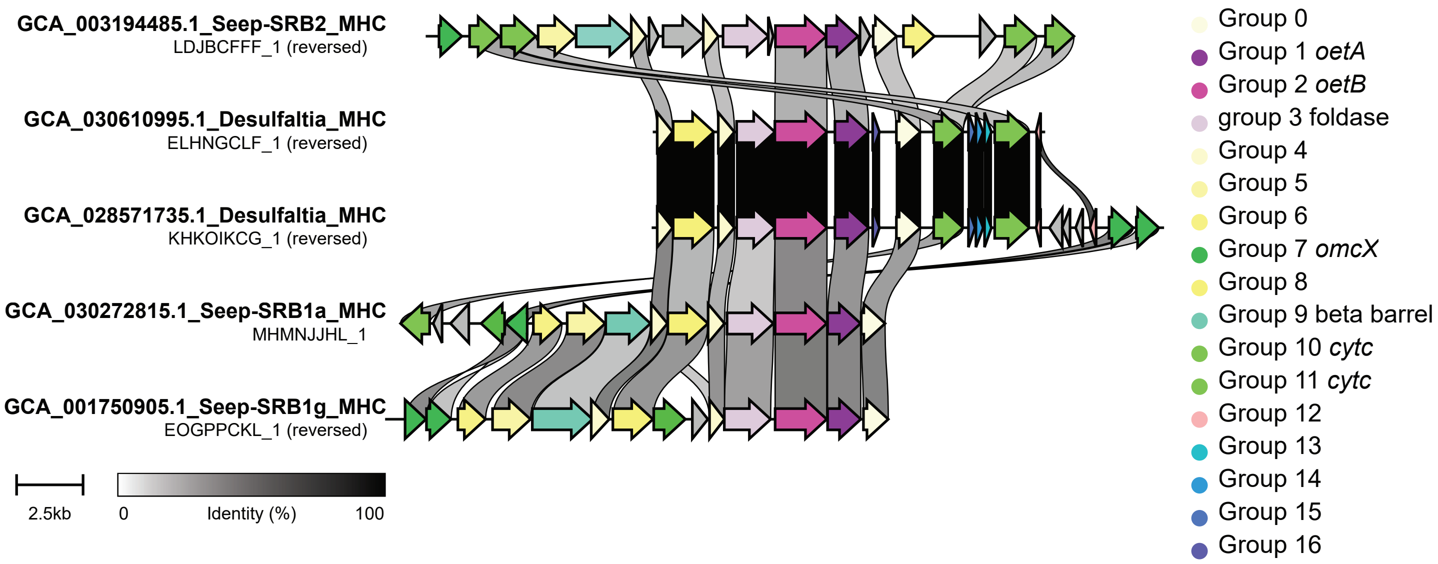


**Supplementary Figure S6 Multiheme cytochrome gene cluster with *oetA* and *oetB* of two *Ca*. Desulfaltia MAGs from gtdb in comparison with the gene clusters of representatives of known ANME partner bacteria Seep-SRB1a, Seep-SRB1g and Seep-SRB2.** The *Ca*. Desulfaltia MAG recovered in this study did not contain the MHC gene cluster. If it truly lacks the MHC cluster, either only some Desulfaltia are ANME partners or interspecies interaction works differently. Because MAGs are inherently incomplete there is the possibility that the MHC cluster was not binned or assembled and therefore lacking. *OetA* and *OetB* are arranged as *OetAB* as in other Seep-SRB groups. In green we depict additional cytochromes, including *omcX*, which are not in the same order between MAGs, but usually present close to *OetAB*. The beta barrel was missing in *Ca*. Desulfaltia MAGs, the assembly broke before the expected position. The gene similarity is highest between the two *Ca*. Desulfaltia MAGs. The genes are generally more similar to Seep-SRB1a than Seep-SRB2 in line with *Ca*. Desulfaltia being in the same family as Seep-SRB1a (ETH-SRB1). Further, some genes without annotation are conserved between Seep-SRB groups. The gene cluster comparison was done with clinker.

**Supplementary Figure S7**

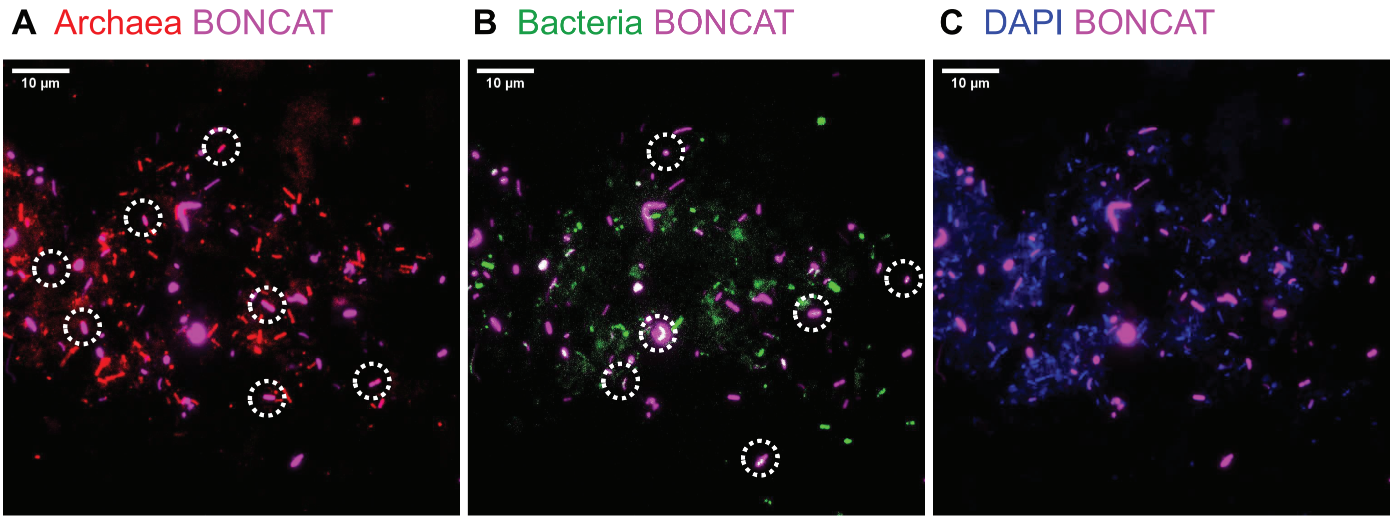


**Supplementary Figure S7 Confirmation of BONCAT active ANME-1 presence in Rock 9.** Here we confirm ANME-1 are a dominant fraction of the community showing their typical rectangular shape. Example images of **A**) ANME-1 cells in red using an ANME-1 probe mix (ANME1-350 and ANME1-728) for a stronger signal, and BONCAT signal indicating translational activity in magenta. For counting we used the brighter archaeal probe (ARCH915). **B**) Bacterial probe mix in green with the BONCAT signal and **C**) DAPI signal in blue and BONCAT signal. White circles highlight examples of respective BONCAT active cells.

**Supplementary Figure S8**

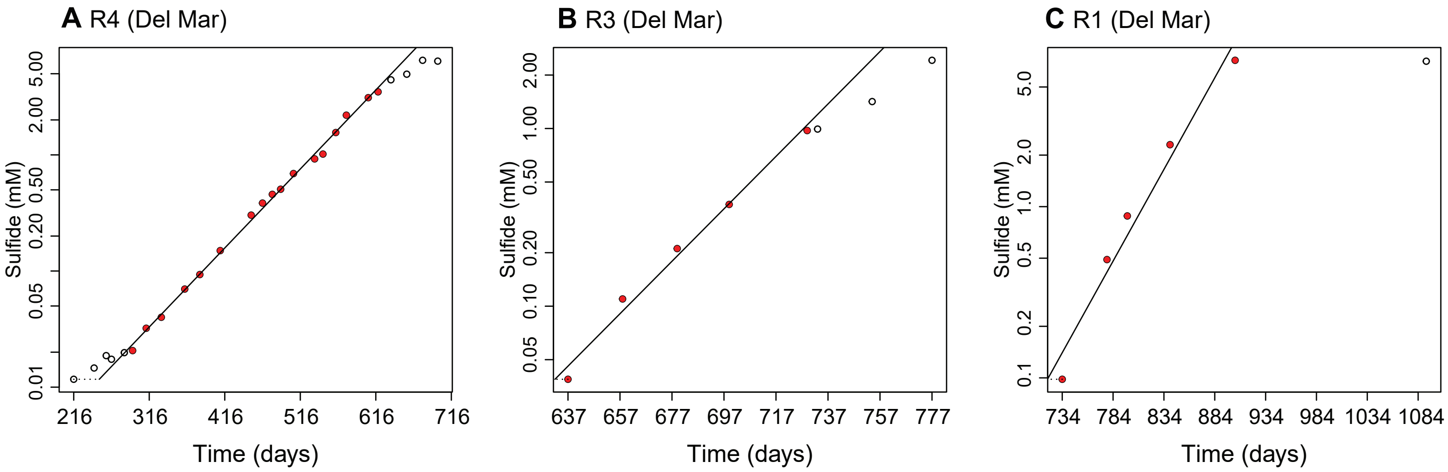


**Supplementary Figure S8 Exponential sulfide increase in long-term incubation of low AOM carbonates with sulfate and methane.** We reactivated the initially low AOM activity carbonates **A**) R4, **B**) R3 and **C**) R1 in order of reactivation, in a long-term incubation mimicking methane resurge with artificial seawater (10 mM sulfate) and a methane headspace under anoxic conditions. The red dots were included in the growth rate calculation. Note that only the phase with increasing sulfide concentrations is shown and therefore not starting at day 0. $R^2$ and growth rate r were derived from a linear regression of log transformed concentrations. The figures show sulfide on a log transformed y-axis. R1 and R3 have even faster doubling times than R4, but compared to R4 less datapoints and lower $R^2$. The long-term incubation of R2 was lost due to accidental breakage of the glass bottle.

### Supplementary Figure S9

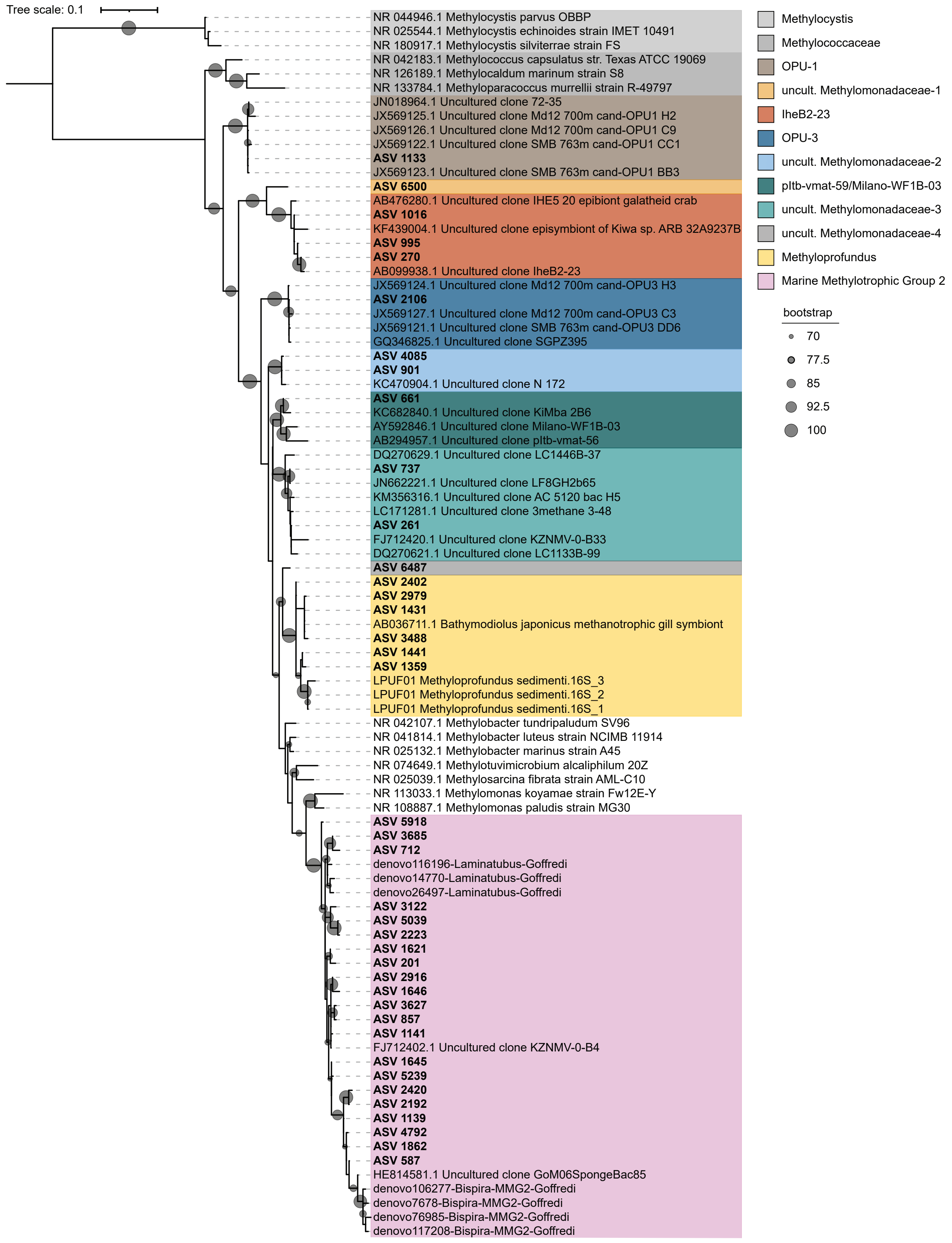


**Supplementary Figure S9 16S rRNA phylogenetic tree of Methylococcales ASVs found in this study with reference sequences following the SILVA taxonomy.** 16S rRNA sequences were first classified with SILVA and Methylococcales were selected. We did not find other known aerobic methanotrophic groups within e.g. Alphaproteobacteria or else. We further classified the Methylococcales sequences according to this tree. For this study, sequences that did not clearly cluster with known groups were named uncultivated Methylomonadaceae-1 through 4. Here we combined pltb-vmat-59 and Milano-WF1B-03 into pltb-vmat-59/Milano-WF1B-03, because they could not be separated based on this tree.

### Supplementary Figure S10

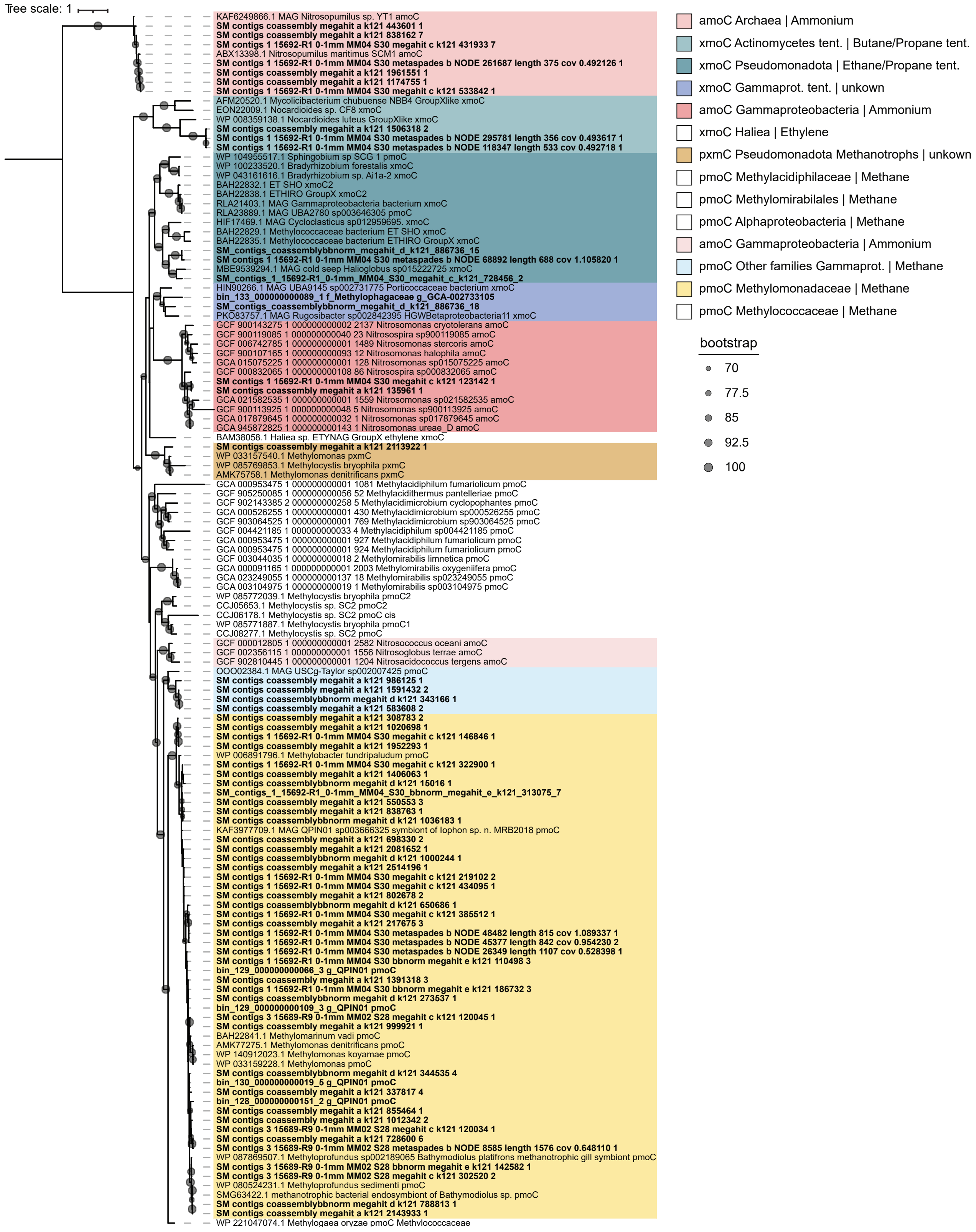


**Supplementary Figure S10 Uncollapsed phylogenetic tree of active site containing subunit C of the CuMMO (*pmoC*/*xmoC*) based on metagenomics.** Sequences in the tree from our study (bold) were assembled into contigs with unknown taxonomic affiliation or are part of a MAG (bin). The clades for ammonium and methane CuMMO are well supported by experimental data in the literature. Although e.g. Mycolicibacterium (synonym Mycobacterium) has been shown to oxidize butane, and ETHIRO (unpublished, isolate lost) has been suggested to oxidize ethane, the diversity within these groups is large, and especially for the seep sequences the substrates represent hypotheses. The *xmoC* Gammaprot. tent. (darker blue) has unknown substrate so far but represents the first CuMMO within Methylophagaceae.
