## Supplementary Table 1 for "Distinct Microbial Communities Within and On Seep Carbonates Support Long-term Anaerobic Oxidation of Methane and Novel pMMO Diversity"

| | | | $^{13}C-CH_4$ | | | $CH_3D$ | | | | $SO_4^{-2}$ |
|---|---|---|---|---|---|---|---|---|---|---|
| Activity | Site | Rock | $^{13}C$-DIC [nmol $cm^{-3}$ $d^{-1}$] | SE | $^{13}C-CH_4$ [%] | Deuterium [nmol $cm^{-3}$ $d^{-1}$] | SE | $CH_3D$ [%] | incubation time (days) | Sulfide |
| low activity | Del Mar | R1 outside | b.d. | | 100 | b.d. | | 100 | 143 | b.d. |
| low activity | Del Mar | R1 inside | b.d. | | 100 | b.d. | | 100 | 140 | b.d. |
| low activity | Del Mar | R2 | b.d. | | 100 | 9.8 | * 1.1 | 100 | 124 | b.d. |
| low activity | Del Mar | R3 | b.d. | | 100 | 51.6 | * 3.2 | 100 | 123 | b.d. |
| low activity | Del Mar | R4 | b.d. | | 100 | 11.6 | 1.6 | 100 | 123 | b.d. |
| interm. | Del Mar | R9.1 (grey) | 106.4 | 4.6 | 100 | 434.2 | 6.6 | 100 | 112 | ✓ |
| interm. | Del Mar | R9.2 (dark grey) | 5.1 | * 0.5 | 100 | 30.9 | 0.4 | 100 | 112 | b.d. |
| high activity | SMM800 | R1 chimlet top | a.d. | | 100 | 3213.3 | 113.4 | 50 | 48 | ✓ |
| high activity | SMM800 | R1 chimlet middle | a.d. | | 100 | 935.7 | 24.0 | 50 | 48 | ✓ |
| high activity | SMM800 | R1 chimlet bottom | a.d. | | 100 | 843.1 | 10.1 | 50 | 48 | ✓ |
| high activity | SMM800 | R3 protochim outside | 2331.1 | 35.8 | 10 | 1070.5 | 22.5 | 50 | 86 | ✓ |
| high activity | SMM800 | R3 protochim inside | b.d. | | 10 | b.d. | | 50 | 86 | b.d. |

b.d. below detection

a.d. above detection

*non-linear

**Supplementary Table 1 Anaerobic methane oxidation activity rates of methane seep carbonates. The first row shows the respective substrates.** Anaerobic methane oxidation was measured with $^{13}C-CH_4$ according to $^{13}C-CH_4 + SO_4^{2-}$ -> $^{13}C-HCO_3^- + HS^- + H_2O$. The labeling percentage added in % is given and was taken into account in the calculation. The incubations with $^{13}C-CH_4$ did not yield a quantitative rate measurement for most samples, due to a combination of an unusually high manufacturer contamination of the $^{13}C-CH_4$ gas with $^{13}C-CO_2$ lowering the sensitivity, low AOM activity or too high activity rendering $^{13}C-HCO_3^-$ production below or above detection. Nevertheless, chimlet with high $CH_3D$ rates was above detection in the $^{13}C-CH_4$ measurements too and even though not quantitative generally agreeing with the $CH_3D$ measurements. Especially for carbonate rocks uncontaminated $^{13}C-CH_4$ is critical. We measured anaerobic methane activation rates (nmol D $cm^{-3}$ $d^{-1}$) in anoxic incubations with monodeuterated methane based on: $CH_3D + SO_4^{2-}$ -> $HCO_3^-$ + $HS^-$ + HDO. We measured water δD over four timepoints and calculate the rate from the linear increase unless stated otherwise. Although the quantitative rates from $^{13}C-CH_4$ and $CH_3D$ are similar, they are not the same. The two isotopes of methane are not measuring exactly the same process. Additional variability might be coming from the rock subsamples as the incubations were done separately on different rock subsamples. The standard error (SE) of k was calculated from the linear regression of the timepoints. The total incubation times (t4) apply to all substrates, with the exception of R9.1 $^{13}C-CH_4$ which was incubated for the same time but the rate was calculated from t0-t3 (day 20), since t4 was above detection. At the last time point (t4) sulfide was detected in R9.1, chimlet top, middle, bottom, and protochimney surface in both $CH_3D$ and $^{13}C-CH_4$ incubations. *Deuterium above background was only detected at t4 indicating a nonlinear increase in R2 and R3, b.d. below detection, surf. surface, int. interior, btm. bottom
