## Supplementary Table 4 for "Distinct Microbial Communities Within and On Seep Carbonates Support Long-term Anaerobic Oxidation of Methane and Novel pMMO Diversity"

| Method | Site and sample type | Doubling time (months) | Growth rate (day$^{-1}$) | Study |
|---|---|---|---|---|
| FISH | Eel River Basin, seep sediment | 8.0 | 0.0028 | Orphan et al.(2009) |
| FISH | Hydrate Ridge, seep sediment | 7.0 | 0.003 | Nauhaus et al. (2007) |
| Bulk sediment $^{15}N$ | Hydrate Ridge, seep sediment | 6.2 | 0.0037 | Kruger et al. (2008)[a] |
| SIMS $^{15}N$ | Eel River Basin, seep sediment | 3.7 (3.0-5.4) | 0.0062 (0.004-0.008) | Orphan et al.(2009) |
| qPCR | Monterey Canyon, seep periphery sediment | 0.7-1 | 0.024-0.031 | Girguis et al. (2005) |
| sulfide | Sediment free enrichment, 60°C | 2.3 | 0.01 | Wegener et al.(2015) |
| nanoSIMS $^{15}N$ | Hydrate Ridge, seep carbonate | 9.0-13.5 | 0.0017-0.0026 | Marlow et al.(2014)[b] |
| nanoSIMS $^{15}N$ | Costa Rica, sediment nodule | 4.5 | 0.01 | Marlow et al.(2014)[b] |
| sulfide | Del Mar, seep periphery carbonate | 1.5 | 0.016 | This study |

a.recalc. Orphan et al. (2009)

b.calc. this study

**Supplementary Table 4 Comparison of the ANME-SRB growth rate measured in our study with ANME-SRB growth rates in the literature.** Our measurement of 44 days (growth rate with highest confidence) was measured in anaerobic long-term incubations with initially low AOM activity carbonates, that were amended with methane and artificial seawater. Sulfide production from AOM was used to calculate the growth rate. Previous ANME-SRB growth rates were measured with different methods including sulfide production (column 1).
